# Serum proteomic atlas reveals distinct molecular signatures of lupus nephritis activity, chronicity, and treatment response

**DOI:** 10.64898/2026.05.08.723858

**Authors:** Rufei Lu, Catriona A. Wagner, Andrea Fava, Ben Jones, Peter Izmirly, H. Michael Belmont, Robert M. Clancy, Jennifer Anolik, Jennifer L. Barnas, Chaim Putterman, David Wofsy, Michael H Weisman, Anne Davidson, Derek M. Fine, V. Michael Holers, Paul J. Utz, the Accelerating Medicines Partnership in RA/SLE Network, Betty Diamond, Jill Buyon, Michelle Petri, Joel M. Guthridge, Judith A. James

## Abstract

Lupus nephritis (LN), a severe manifestation of systemic lupus erythematosus (SLE), features heterogeneous renal pathology and reliance on invasive biopsies for diagnosis, prognosis, and treatment selection. Current peripheral clinical markers inadequately capture disease activity and progression. Here, we performed comprehensive serum proteomic profiling of over 5,000 proteins in the large, longitudinal Accelerating Medicines Partnership Rheumatoid Arthritis/SLE cohort of 270 LN patients and 63 healthy controls. Machine learning identified distinct molecular signatures that classified LN versus controls, differentiated histological classes, and delineated activity- and chronicity-associated pathways, including inflammatory cytokine, PI3K/AKT, TGFb, and complement/coagulation pathways. An increase in VSIG4, CD27, HAVCR1, and LAIR1 consistently emerged as top biomarkers across multiple clinical contexts, and early decreases in these markers at 3 months were associated with complete treatment response at 1 year. By resolving coordinated serum protein modules linked to key inflammatory, PI3K/AKT, TGFb, and complement pathways, these signatures mechanistically connect circulating proteomic perturbations to intrarenal immune activation, tissue injury, and repair in LN. These findings demonstrate that serum proteomics reflect complex intrarenal immunopathology and offer a promising noninvasive “liquid biopsy” approach to refine LN classification and guide personalized management, potentially reducing the need for repeated invasive biopsies and improving therapeutic decision-making.

## Introduction

Lupus nephritis (LN), a major manifestation of systemic lupus erythematosus (SLE), affects up to half of patients with SLE and is a leading cause of morbidity and early mortality (1). Clinical decision-making in LN relies on invasive renal biopsy for diagnosis and histologic classification, as well as on clinical markers, such as proteinuria, serum creatinine, and complement levels. The International Society of Nephrology/Renal Pathology Society (ISN/RPS) system stratifies LN into 6 classes based on patterns of glomerular inflammation and chronic injury, with proliferative classes (class III and IV, with or without concurrent class V membranous features) guiding the need for intensive immunosuppression (2–4). In addition, the National Institutes of Health Activity Index (NIH-AI) and Chronicity Index (NIH-CI) are histopathological scoring systems that further quantify the degree of renal inflammation and fibrosis, respectively, with higher activity scores informing treatment decisions and prognosis (5). These indices reflect a biological continuum in which LN progresses from an active, inflammatory phase to a chronic phase, with persistent immune activation leading to tissue remodeling and permanent fibrotic scarring.

Despite these assessment tools, significant challenges remain in LN management. Kidney histopathology can evolve, with more than half of patients switching histologic class on repeat biopsy, often necessitating changes in treatment (6). In addition, persistent elevation of NIH-AI at repeat biopsy predicts subsequent renal flares and poorer long-term survival, even when proteinuria has resolved (7, 8). These findings underscore the value of serial kidney evaluation; however, repeat biopsies are invasive and impractical for routine longitudinal monitoring. Although clinical biomarkers could reduce the need for repeated tissue sampling, currently available measures are imperfect and often discordant with histologic activity, with up to one-third of patients exhibiting ongoing renal inflammation despite clinical quiescence (9, 10). Thus, there is an urgent need for noninvasive biomarkers that accurately reflect the underlying renal histology of LN.

Furthermore, despite advances in therapy, current LN outcomes remain suboptimal, with up to 30% of patients still progressing to end-stage kidney disease within 10 years (11). Even among those classified with proliferative disease, response to therapy varies markedly. Notably, the pivotal trials of belimumab and voclosporin enrolled only patients with ISN/RPS class III, IV, or V, yet nearly 60% failed to achieve their primary endpoints (12, 13), underscoring that substantial biological heterogeneity exists even within these histologically similar groups. Thus, although clinically informative, current classification systems insufficiently capture the molecular diversity that drives individual disease trajectories. Together, these limitations highlight the need for molecularly informed, noninvasive biomarkers that capture intrarenal immunopathology and predict treatment response and outcome.

While urinary biomarkers have emerged as valuable tools for assessing local renal injury, longitudinally collected serum samples represent an ideal complementary biospecimen for systemic inflammation. Serum is noninvasive and clinically accessible, even in patients with limited renal output, and may reflect dynamic changes in systemic immune responses that drive local tissue pathology. Prior studies have identified promising associations between select serum and plasma soluble biomarkers, such as TWEAK, ALCAM, syndecan-1, VCAM-1, and anti-C1q, and histologic activity or prognosis in LN (14–21). More recently, high-dimensional proteomic studies in urine have quantified up to ∼1,000 proteins and linked specific signatures, including macrophage activation markers such as CD163, to proliferative histology, NIH activity and chronicity indices, and treatment response (22–26). However, large-scale proteomic profiling of serum in biopsy-confirmed LN remains limited, and existing serum studies typically investigate far smaller biased panels, thereby limiting their ability to capture the extensive molecular heterogeneity of LN. Because disease pathways and clinical responses are highly variable across individuals and over time, relying on a restricted panel of biomarkers may overlook key biological processes. To address this gap, our study comprehensively profiled more than 5,000 serum proteins, enabling a robust, unbiased approach to molecular discovery, LN classification refinement, and treatment outcome prediction.

## Results

### Preliminary identification of protein biomarkers for lupus nephritis

To delineate the serum proteomic landscape of LN, we first profiled 1,536 proteins in a preliminary, independent cohort of SLE patients with ACR-defined renal involvement (n=27) and age- and sex-matched HCs (n=16). All participants were female, with comparable median ages (LN: 29 years [IQR 24-39]; HCs: 30.5 years [IQR 24-41]; Supplemental Table 1). The distribution of race did not differ significantly between LN patients and HCs, but the proportion of Hispanic or Latino individuals was significantly higher in the LN group (41%) than in HCs (6%; Supplemental Table 1). Unsupervised UMAP clustering of all measured proteins demonstrated a distinct separation between patients with renal involvement and HCs (Supplemental Figure 1a), indicating profound systemic proteomic changes in LN. Supervised machine learning (iterativew XGBoost) achieved high classification accuracy (87.9% ± 0.1%; 10-fold cross-validation) and robust discrimination (AUC = 0.96 ± 0.07; Supplemental Figure 1b). A total of 400 proteins showed significantly different abundance in LN compared to HCs (273 increased; 127 decreased; Supplemental Data 1). Of these, the most important predictors, all increased in LN, included HAVCR2, BST2, TNFRSF1B, CD27, VSIG4, CD300C, CD300E, VCAM1, TNFRSF8, and CXCL9 (Supplemental Figure 1c-d and Supplemental Table 2).

### Validation and expansion of LN protein biomarkers in the AMP SLE longitudinal cohort

Building on these preliminary findings, we expanded our analysis to an independent longitudinal AMP cohort of 270 patients with biopsy-confirmed LN and 63 HCs. This cohort was predominantly female (85%) with a median age of 34 years, similar to HCs (Table 1). Most LN patients self-identified as Black (n=118; 44%) or White (n=76; 28%; Table 1). The cohort spanned the full spectrum of ISN/RPS LN classes, including 93 (34%) pure proliferative (ISN/RPS class III/IV), 78 (29%) mixed proliferative (class III/IV + V), 64 (24%) membranous (ISN/RPS class V), 23 (9%) mesangial (ISN/RPS class I/II), and 12 (4%) advanced sclerosing LN (Table 1). Approximately one-third of LN patients were undergoing their first renal biopsy at study entry. LN patients frequently exhibited positivity for anti-dsDNA antibodies (67%) and hypocomplementemia (59% low C3; 51% low C4). Patients had evidence of systemic disease activity (mean hybrid SELENA-SLEDAI score, 12.2; SD, 6.0), with 1/3 demonstrating extrarenal manifestations (excluding complement or anti-dsDNA activity alone). Measures of renal involvement included a mean UPCR of 2.9 (SD, 2.6), a mean estimated glomerular filtration rate of 69 mL/min/1.73m^2^ (SD, 31.4), and a mean serum creatinine of 1.2 mg/dL (SD, 0.82), reflecting moderate renal impairment (Table 1).

**Table 1.**
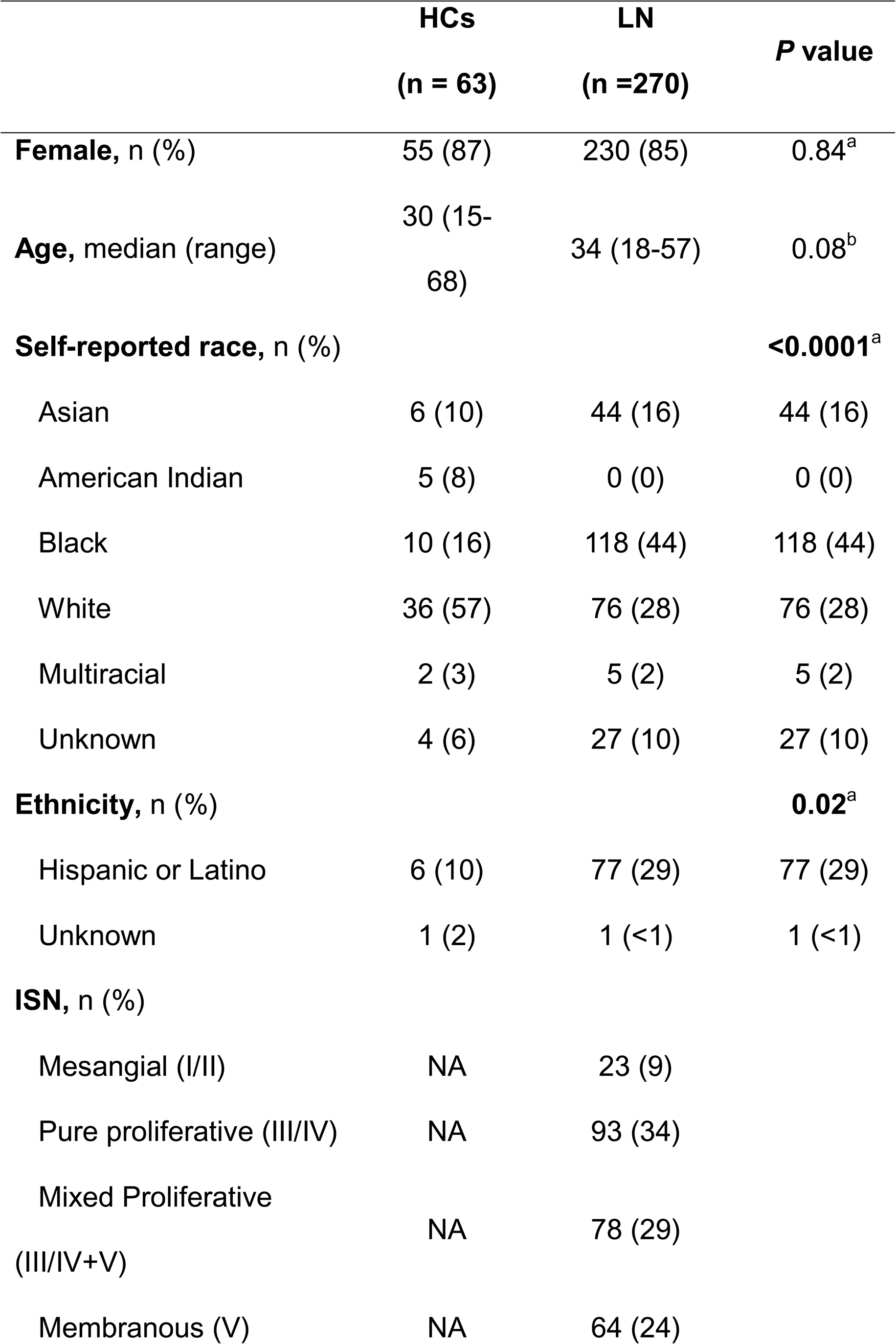
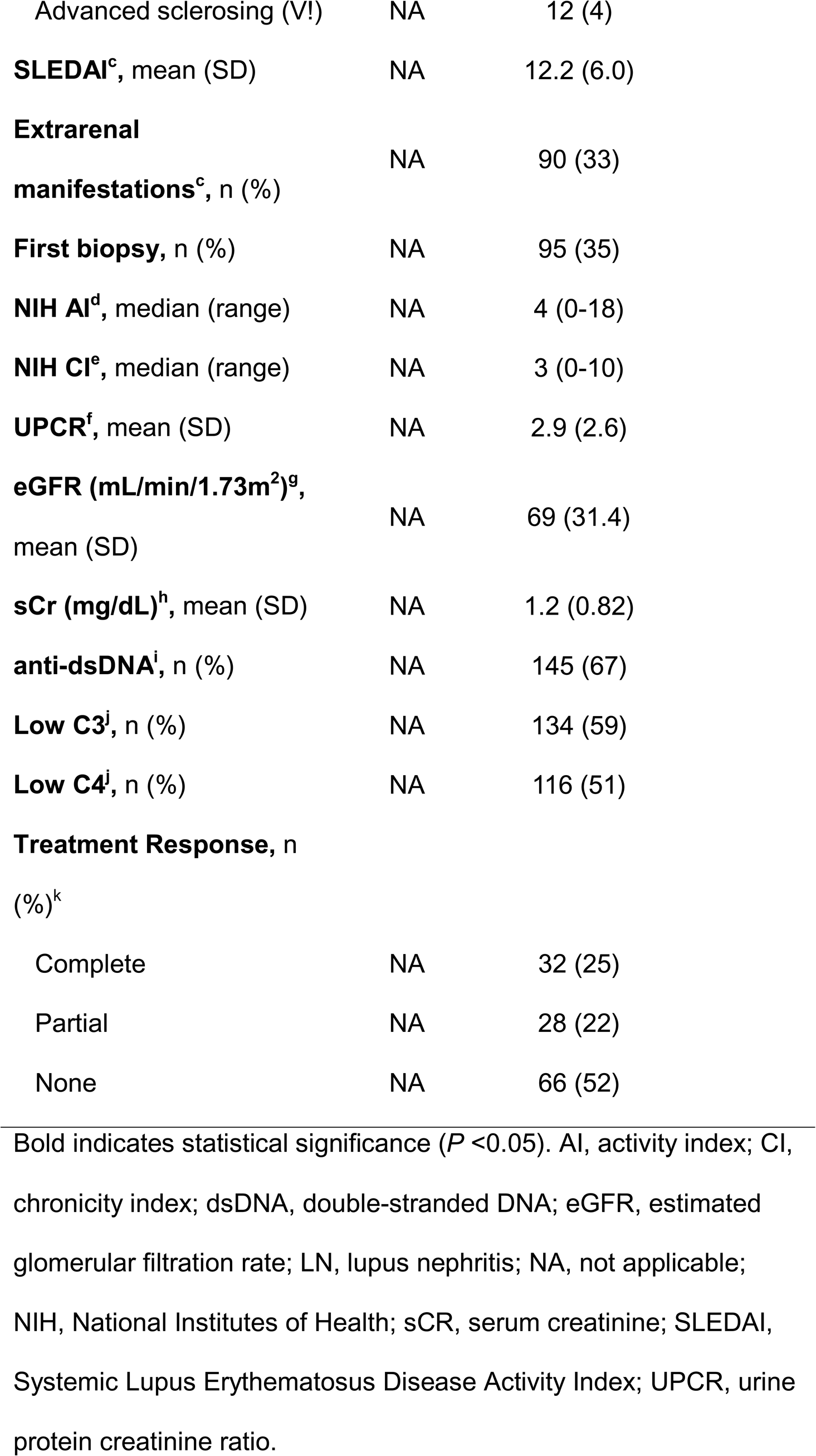
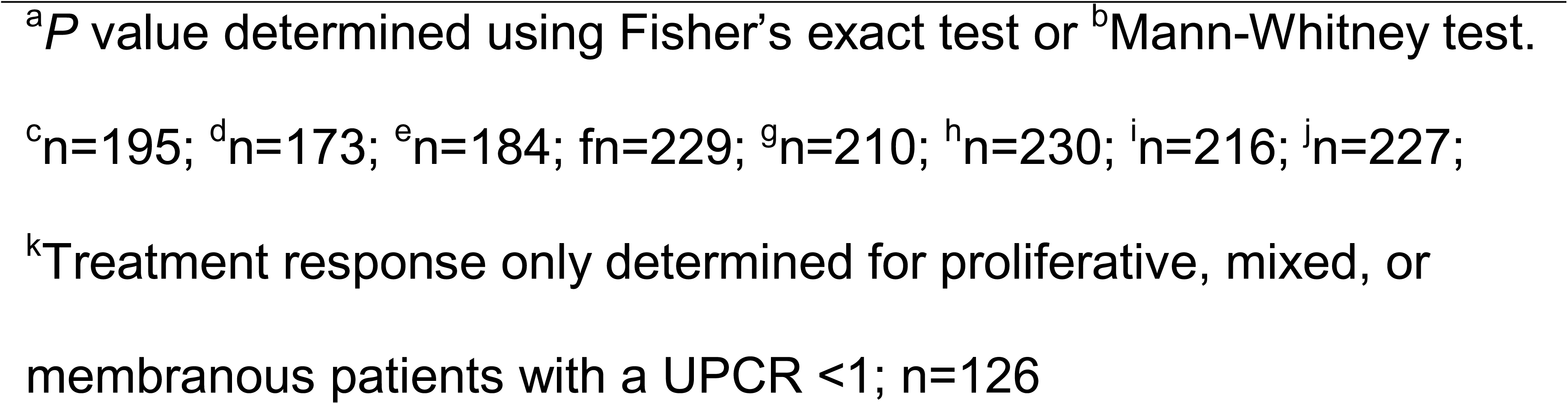
Demographics and clinical characteristics of patients with lupus nephritis (LN)

Proteomic profiling quantified a final set of 5,095 proteins per sample, following a rigorous multi-step QC pipeline (Figure 1a and Supplemental Figure 2a). Consistent with the preliminary cohort, UMAP visualization revealed robust separation between LN patients and HCs across all baseline and longitudinal samples (Figure 1b and Supplemental Figure 2b). Differential expression analysis revealed extensive soluble proteomic dysregulation in LN patients compared to HCs, with 2,539 proteins significantly increased and 371 decreased in LN (Supplemental Data 2 and Supplemental Figure 2c). Through an iterative boosted decision tree machine learning algorithm, we identified a panel of 41 proteins that distinguished SLE patients with LN from HCs with high accuracy (97% ± 2%) and discrimination (AUC = 0.99 ± 0.01; Figure 1c-d and Supplemental Table 3). The most important predictors included increased levels of TNFRSF1B, VSIG4, HAVCR2, CD27, and BST2, all of which were also identified in the preliminary cohort (Figure 1c, Supplemental Figure 1c-d, and Supplemental Table 3).

**Figure 1.**
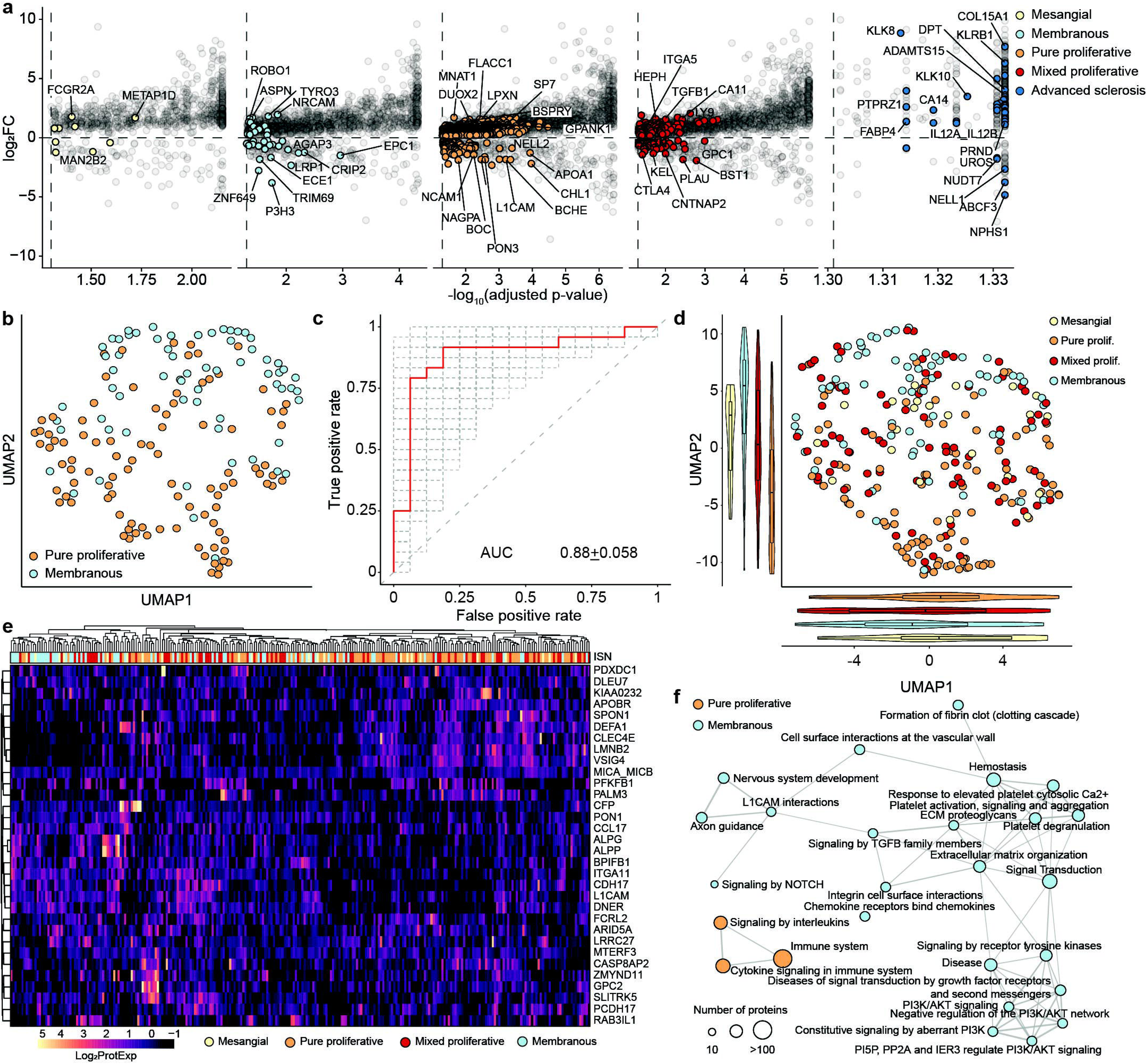
Baseline serum proteomic profiling distinguishes lupus nephritis patients from healthy controls. **(a)** Study design showing enrollment of 270 SLE patients with biopsy-confirmed lupus nephritis (LN) and 63 healthy controls (HC). Serum samples were collected at baseline (time of biopsy), 3, 6, and 12 months. ISN class and NIH activity/chronicity indices were determined from renal biopsies, and treatment response was assessed at 12 months. Serum samples were analyzed using the Olink Explore HT ® platform for 5,095 high-quality proteins. **(b)** UMAP visualization of serum proteomic profiles from LN patients and HCs based on the 5,095 proteins. **(c)** Volcano plot displaying quantified proteins according to log2 fold change and importance values for classifying LN and HCs. Dashed lines indicate the importance threshold (elbow point) for selecting top predictors (n=41) and fold-change cut-offs (±0.2). **(d)** ROC curve demonstrating the classification performance of the 41-protein biomarker panel for distinguishing LN from healthy controls, evaluated by 500-fold cross-validation. Gray dashed lines represent the ROC curves from individual cross-validation iterations, while the red curve denotes the average ROC across all iterations. **(e)** Heatmap of significantly enriched KEGG pathways in LN compared to HCs. **(f)** Network visualization of enriched REACTOME pathways, with node size representing the number of proteins and line thickness indicating pathway connectivity. **(g-j)** Volcano plots showing protein enrichment from **(g)** KEGG TLR signaling pathway, **(h)** REACTOME complement cascade, and **(i)** REACTOME interferon signaling pathways. Dashed lines indicate significance thresholds (*P_adj_* = 0.05) and fold-change cut-offs (±0.2). **(j)** LN patients were stratified based on exhibiting only renal SLEDAI domains (Renal-Limited; n=105) or renal concurrent with extrarenal domains, excluding anti-dsDNA and complement (Renal-Plus; n=90). The scatter plot displays the log2FC relative to HCs for all proteins identified as differentially expressed (Padj < 0.05) in either analysis. Points are stratified by significance overlap (shape) and colored by immune pathway classification (innate vs. adaptive) using the REACTOME database. Statistical significance was determined using logistic regression, adjusting for covariates including age, sex, and genetic ancestry.

To understand the biological context of the differentially detected proteins (DEPs), we performed pathway enrichment analyses using GO, KEGG, and REACTOME databases, revealing a robust increase of immune activation processes in LN, particularly cytokine signaling (IFN, TNF, TLR), adaptive immune responses (Th1/Th2 cell differentiation, antigen presentation), and complement activation (Figure 1e-f and Supplemental Figure 3a). We further validated these trends by examining individual proteins, confirming elevated expression of key mediators in TLR signaling (STAT2, MAP2K6), complement pathways (C3, C4B), and IFN signaling (GBP4, TRIM25, STAT2; Figure 1g-i). Additionally, disease-associated changes in neurodegeneration (PPP3R1, BDNF, NEFL, GFAP), neutrophil degranulation (TNFRSF1B, LAMP1), and platelet activation (GP1BA) proteins further underscore the broad dysregulation of immunity and cellular structure in LN (Supplemental Figure 3b-d).

To distinguish proteomic alterations driven specifically by renal pathology from those reflecting broader systemic involvement, we stratified LN patients by SLEDAI score into “Renal-Limited” (active renal domains only) and “Renal-Plus” subgroups (active renal domains with concurrent extrarenal manifestations), excluding anti-dsDNA and complement, which could be present in either subgroup. Comparing these subgroups to HCs confirmed robust dysregulation in both phenotypes, with 1,312 increased and 268 decreased proteins in Renal-Limited patients, and 1,724 increased and 253 decreased proteins in Renal-Plus patients (Supplemental Data 3). Notably, 996 (63% of those in Renal-Limited; 50% of those in Renal-Plus) of these proteins were concordantly altered in both subgroups, highlighting a shared systemic proteomic signature (Figure 1j). Direct comparison between the two clinical phenotypes revealed 217 DEPs; 187 proteins were elevated in the Renal-Limited group, including C1QA, VCAM1, HAVCR2, and VSIG4, while only 30 were higher in the Renal-Plus group (Supplemental Data 3 and Figure 1j, suggesting that these proteins may reflect heightened renal-specific pathology rather than the broader systemic inflammation observed in patients with broader organ involvement.

### Proteomic signatures of LN histological class

We next investigated whether specific soluble protein signatures could further resolve the molecular heterogeneity across LN histological classes. DEP analysis revealed varying degrees of dysregulation across ISN classes when compared to HCs (Supplemental Data 4). Pure proliferative LN exhibited the most extensive differences with 3,139 DEPs, followed by mixed proliferative (2,383 DEPs), membranous (1,706 DEPs), and mesangial (975 DEPs). Notably, advanced sclerosing LN showed minimal dysregulation, with only 578 DEPs, suggesting a shift toward cellular loss and a quiescent/fibrotic phase. To identify class-defining features, we highlighted proteins uniquely enriched in each histological class (Figure 2a). Pure and mixed proliferative LN exhibited the most distinct molecular profiles compared to age-, sex-, and genetic ancestry-matched HCs, while mesangial LN appeared the least distinct proteomic signature compared to the other LN classes (Figure 2a). Despite these unique class-specific markers, substantial molecular overlap exists between classes (Supplemental Figure 4). For example, 1618 DEPs (52% of pure proliferative DEPs; 95% of membranous DEPs) were shared between pure proliferative and membranous LN, including those involved in cytokine-receptor interaction (TNFRSF1A, IFNAR2; Supplemental Figure 4a). Incorporation of mixed proliferative LN further increased the overlap, with 2,190 DEPs shared between pure proliferative and mixed proliferative LN (70% of pure proliferative; 92% of mixed proliferative), and 1,572 shared between membranous and mixed proliferative LN (92% of membranous; 66% of mixed proliferative; Supplemental Figure 4b-c). Mesangial LN showed substantial convergence with both proliferative and membranous classes, with 953 DEPs shared with pure proliferative LN (30% of pure proliferative; 98% of mesangial) and 888 DEPs shared with membranous LN (52% of membranous; 91% of mesangial; Supplemental Figure 4d-e). In contrast, advanced sclerosing LN demonstrated the least overlap, sharing 508 DEPs with pure proliferative LN (16% of pure proliferative; 88% of advanced sclerosing) and 434 DEPs with membranous LN (25% of membranous; 75% of advanced sclerosing; Supplemental Figure 4f-g). Pathway enrichment analyses confirmed both shared and unique biology across histological classes. For example, cytokine-cytokine receptor interactions were commonly dysregulated across pure proliferative, mixed proliferative, membranous, and mesangial LN, while cytokine response pathways were shared between pure and mixed proliferative LN (Supplemental Figure 4c).

**Figure 2.**
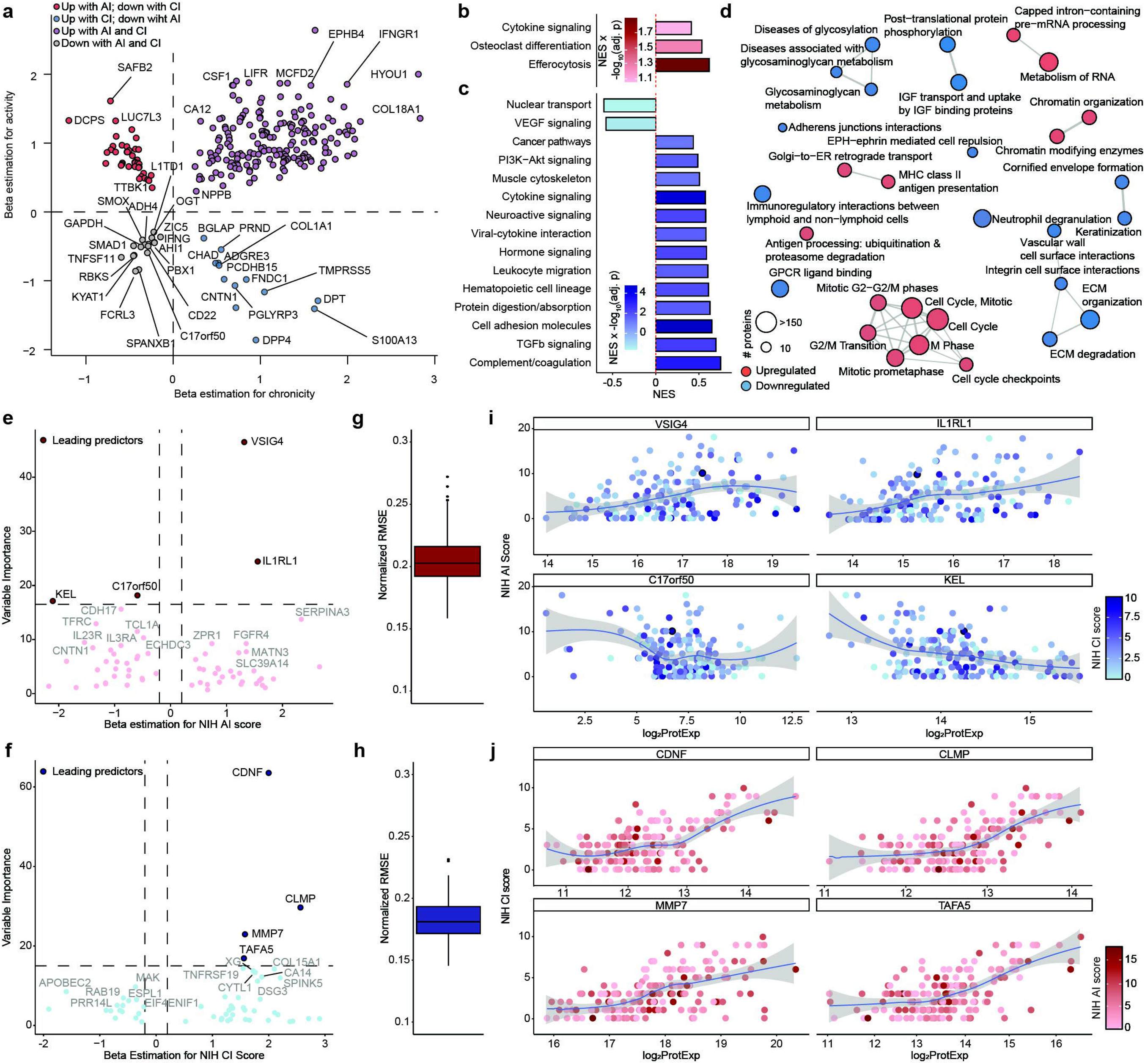
Soluble proteomic profiling defines distinct and shared molecular signatures across lupus nephritis histological classes. Soluble proteomic profiles were determined in patients with pure proliferative (n=93; class III/IV), mixed proliferative (class III/IV + V; n=78), membranous (class V; n=64), mesangial (class I/II; n=23), and advanced sclerosing (class VI; n=12) LN. **(a)** Volcano plots of differentially detected proteins (DEPs) across ISN classes compared to healthy controls. Colored points represent DEPs (*P_adj_* <0.05 and log_2_FC ±0.2) distinct to each class. Gray points indicate shared DEPs. Dashed lines mark significance thresholds. Statistical significance was determined using logistic regression, adjusting for covariates including age, sex, and genetic ancestry. **(b)** UMAP plot derived from 32 discriminatory proteins identified through iterative Boosted Tree Machine Learning analysis for distinguishing membranous and pure proliferative LN patients. Each point represents an individual patient, colored by ISN class. **(c)** ROC curve demonstrating the classification performance of the 32-protein biomarker panel for distinguishing LN from healthy controls, evaluated by 500-fold cross-validation. Gray dashed lines represent the ROC curves from individual cross-validation iterations, while the red curve denotes the average ROC across all iterations. **(d)** Application of the 32-protein model to patients with mixed proliferative and mesangial LN. Points represent individual patients, colored by ISN class. **(e)** Heatmap of the protein expression of the 32 top discriminatory proteins in patients with LN. Protein expression values were log_2_ transformed, and hierarchical clustering was performed using Euclidean distance. **(f)** Significantly different pathways (FDR<0.05) are shown based on enrichment analysis of the top 100 differentially expressed proteins between membranous and pure proliferative LN, as identified by logistic regression models adjusted for age, gender, and genetic ancestry.

To test whether proteomic signatures could distinguish histological classes, we constructed a targeted model using an iterative ensemble boosting approach and identified 32 proteins that robustly discriminate pure proliferative from membranous LN (Supplemental Table 4). UMAP visualization of these biomarkers separated the two classes, achieving 80.9% ± 6.05% classification accuracy and minimal overlap (Figure 2b). ROC analysis confirmed strong discriminatory performance (AUC = 0.88 ± 0.058; Figure 2c). When this model was applied to mixed proliferative and mesangial patients, these classes were found across the proteomic continuum (Figure 2d), a finding supported by heatmap clustering of the 32 discriminant proteins, which demonstrates substantial within-class heterogeneity (Figure 2e). Pathway network analysis of the top 100 discriminatory proteins revealed distinct molecular mechanisms underlying proliferative and membranous LN (Figure 2f). In particular, membranous LN was associated with dysregulation of cellular adhesion, extracellular matrix organization, and PI3K/AKT pathways; pure proliferative LN was associated with predominantly inflammatory response pathways (Figure 2f).

Finally, given the unique molecular signature of advanced sclerosing LN, a dedicated classifier effectively distinguished this group from all other classes (accuracy 90.7% ± 2.1%; AUC 0.92 ± 0.095; Supplemental Figure 5a-b). Key discriminatory proteins included fibrosis markers (COL15A1, MMP10) and tissue remodeling factors (FOLR2, SMAD5), consistent with end-stage renal pathology (Supplemental Figure 5c).

### Proteomic profiles of histological activity and chronicity

We next analyzed serum proteomic profiles underlying histologic activity and chronicity in a subset of patients with proliferative or membranous LN (n=170). Within this cohort, histologic activity and chronicity indices exhibited distinct yet partially overlapping distributions (Supplemental Figure 6). To define proteomic signatures of these pathologic indices, we applied rank-based regression modeling adjusted for age, sex, and genetic ancestry to identify 509 proteins significantly associated with the AI and 2,270 proteins associated with the CI (Supplemental Figure 7 and Supplemental Data 5-6). While the proteomic signatures of activity and chronicity were largely distinct, we identified 253 DEPs (50% of those associated with AI; 11% of those associated with CI) that were shared between the two indices (Figure 3A). Notably, 201 (n=103; 79%) exhibited concordant associations (Figure 3a). Pathway enrichment analyses further supported distinct and partially overlapping mechanisms: increased activity scores were enriched for cytokine-mediated signaling and efferocytosis, whereas chronicity scores were characterized by increased representation of TGFb signaling, complement and coagulation cascades, PI3K-AKT signaling, and leukocyte migration/hematopoietic lineage pathways, together with broader cell cycle and antigen presentation pathways (Figure 3b-d). Importantly, cytokine signaling pathways were also significantly enriched in chronicity scores (Figure 3c), mirroring the shared inflammatory signature observed in the individual protein analysis.

**Figure 3.**
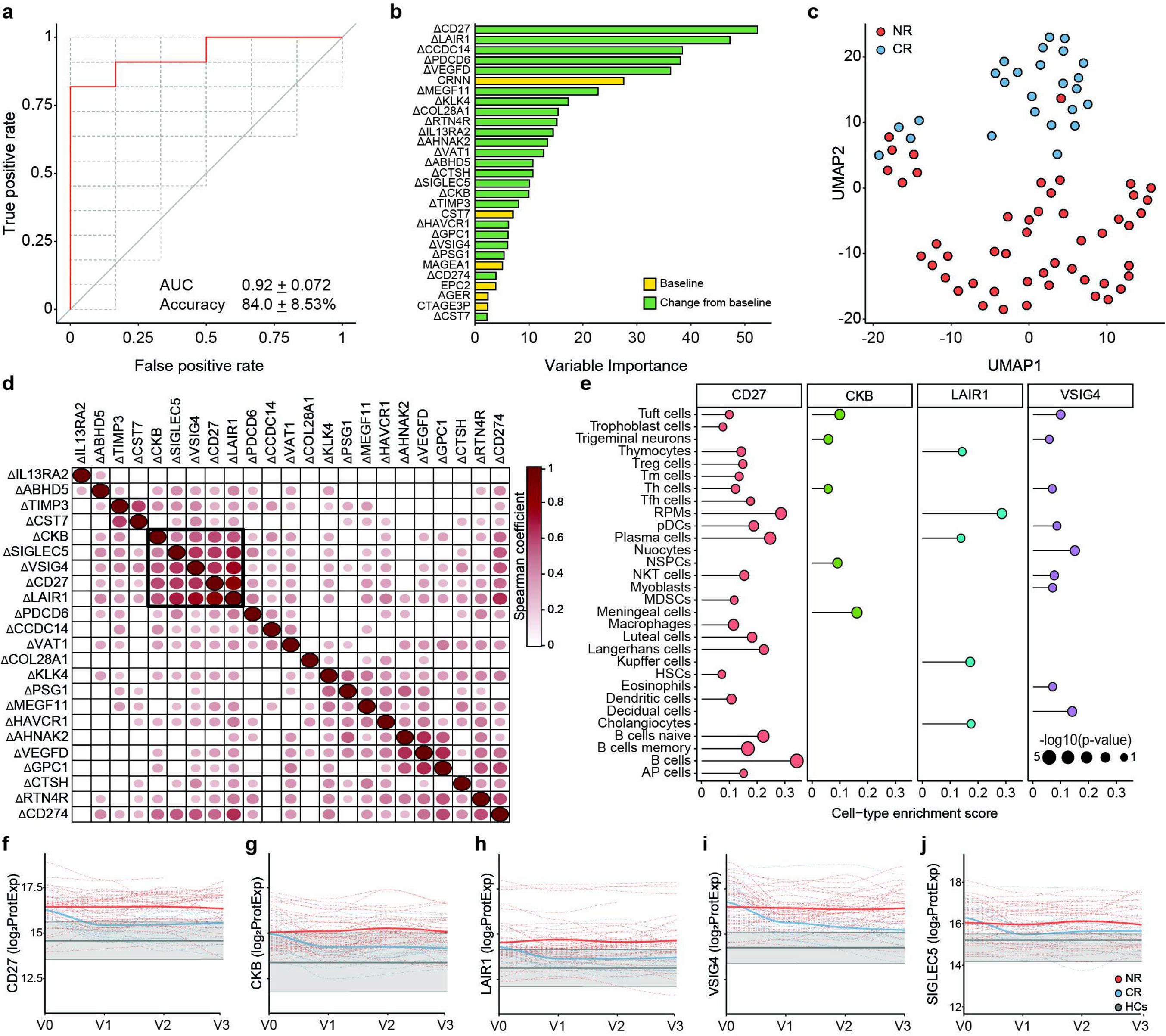
Proteomic profiles associate with NIH activity (AI) and chronicity indices (CI) in patients with proliferative or membranous LN (n=170). **(a)** Bidirectional volcano plot of the beta coefficients for associations between individual proteins and the NIH AI and CI. Only proteins significantly (FDR < 0.05) associated with both AI and CI are shown. Statistical significance was determined using non-parametric regression, adjusting for covariates including age, sex, and genetic ancestry. **(b-c)** Gene set enrichment analysis (GSEA) of proteomic predictors with **(b)** NIH AI and **(c)** NIH CI using the KEGG database. **(d)** Network plot of Reactome pathways significantly associated with NIH CI. **(e-j)** Non-parametric ranked-based regression analyses were performed for **(e, g, i)** NIH AI and **(f, h, j)** NIH CI using an iterative Boosted Tree Machine Learning algorithm. **(e-f)** Volcano plots of variable importance and beta coefficients for the top predictors selected to construct the final model. Within this group, a subset of leading predictors was identified via the elbow method. **(g-h)** Predictive accuracy of the final models, constructed using the full set of top predictors. Box plots indicate the distribution of normalized root mean square error (RMSE) values from predictive models, representing the model’s average deviation from actual scores after scaling to the data range. Lower normalized RMSE values reflect better predictive accuracy. **(i-j)** Regression plots showing the relationship between leading predictor proteins and NIH indices for **(i)** NIH AI with point shading according to individual CI scores and **(j)** NIH CI with point shading according to individual AI scores.

An iterative XGBoost rank-based regression approach identified distinct sets of top linear and non-linear protein correlates for both indices (Figure 3e, f, and Supplemental Tables 5-6). Machine learning regression models constructed from these top proteins demonstrated robust accuracy for both AI and CI, as measured by RMSE (Figures 3g and 3 h). For the NIH AI, the leading predictors included increases in VSIG4 and IL1RL1 and decreases in KEL and C17orf50 (Figure 3e,g). Visualizing these predictors against CI scores confirmed that these associations persist independent of the underlying fibrotic burden (Figure 3e, i). For the NIH CI, the top predictors were increases in CDNF, CLMP, MMP7, and TAFA5; these associations appeared independent of inflammatory activity (Figure 3h,j), indicating that they capture accumulated damage even in the absence of active disease. Together, these analyses underscore the complex proteomic landscape distinguishing local histologic activity from chronic damage.

### Proteomic signatures of treatment response

Having established that serum proteomic signatures can distinguish LN histological classes and correlate with disease activity and chronicity, we next investigated whether these molecular patterns could predict and track therapeutic response. Within our cohort, 34 patients achieved a complete response within 1 year, whereas 66 did not. We first developed a predictive model for complete response using baseline serum protein levels, which achieved moderate discrimination (AUC, 0.8±0.09; Accuracy, 76%±7%), and identified TTLL5, RIN1, CSTT7, VSIG2, YPO10, and BLM as the top predictors (Supplemental Figure 8). As changes in protein levels might better reflect treatment response, we integrated both baseline levels and early changes (from baseline to week 12), which demonstrated substantially stronger discrimination (AUC, 0.92 ± 0.072; Accuracy, 84% ± 8.53%; Figure 4a), driven by a distinct set of proteins enriched for immune regulatory molecules (Figure 4b). UMAP visualization based on these predictors confirmed a separation between complete responders and non-responders (Figure 4c). For reference, a model using only visit 3 protein levels yielded the highest discrimination (AUC = 0.95), though this reflects the post-treatment state at the time of response assessment and therefore lacks prognostic and therapy guidance utility (Supplemental Figure 9). Notably, several proteins associated with NIH-AI and -CI, including VSIG4, CD27, and HAVCR1, consistently emerged as main predictors in both the change-from-baseline and visit 3 models, underscoring their relevance for LN-dominant pathogenesis and differential treatment response (Figure 4b and Supplemental Figure 9).

**Figure 4.**
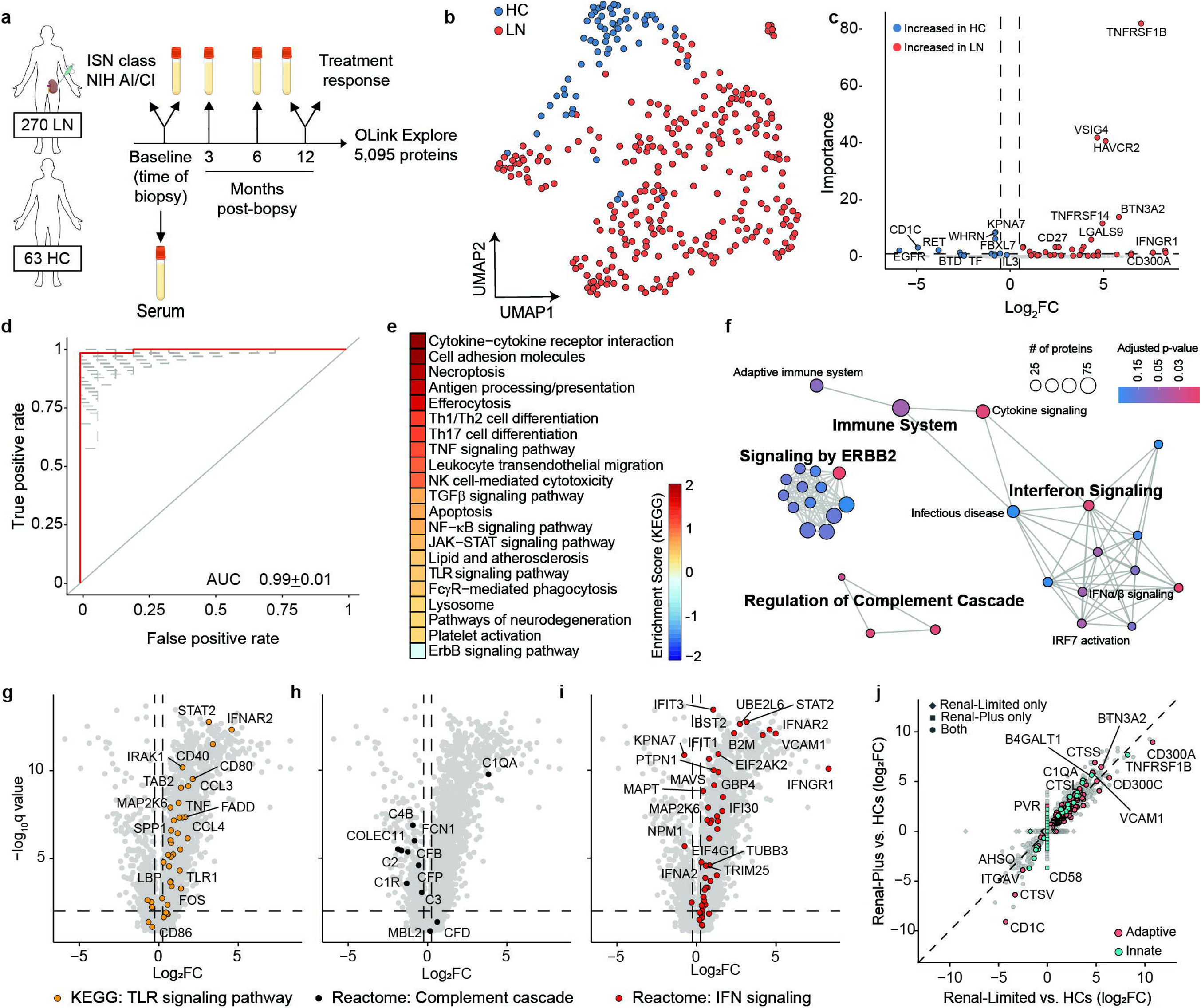
Early changes in serum proteomic profiles predict 1-year treatment response in lupus nephritis (LN) **(a)** ROC curve showing the performance of the iterative Boosted Tree Machine Learning classifier in distinguishing patients with complete response (CR; n=32) from those with no response (NR; n=66) at 1 year. The model was initially trained using all available baseline protein levels and early changes (from baseline to week 3) in serum proteins, and the most discriminatory predictors were selected through an iterative feature elimination process. ROC curves from individual 500-fold cross-validation iterations are shown as gray lines, while the red curve denotes the average performance across all iterations. **(b)** Variable importance plot highlighting the top predictors for treatment response. **(c)** UMAP visualization of serum proteomic profiles for patients with CR and NR, using the top discriminatory proteins. **(d)** Spearman correlation matrix of the top predictors (change from baseline to week 3), illustrating highly correlated protein modules. Hierarchical clustering was performed using Euclidean distance to group proteins with similar correlation profiles. **(e)** Cell-of-origin enrichment analysis for the top four correlated predictors, indicating their predominant cellular sources. **(f-i)** Longitudinal changes in **(f)** CD27, **(g)** CKB, **(h)** LAIR1, **(i)** VSIG4, and **(j)** SIGLEC5 levels over 1 year in patients with CR and NR. Protein expression values are log_2_-transformed. Bold line indicates the mean value. Gray lines indicate the mean and IQR for healthy controls (HCs). Treg, T regulatory; Tm, T memory; Th, T helper; Tfh, T follicular helper; RPMs, red pulp macrophages; pDCs, plasmacytoid dendritic cells; NSPCs, neural stem/precursor cells; NKT, natural killer T cells; MDSCs, myeloid-derived suppressor cells; HSCs, hematopoietic stem cells; AP, anterior pituitary gland

Intra-personal non-parametric correlation analysis of the top change-from-baseline predictors revealed a tightly correlated module comprised of CD27, CKB, LAIR1, VSIG4, and SIGLEC5 (Figure 4d). Network analysis demonstrated that these proteins and their interactors were enriched in immune-related surface receptor binding and signaling pathways (Supplemental Figure 10), while cell-of-origin enrichment suggest that these proteins originate from a range of immune cell types, including B and T cells (CD27) and myeloid cells (LAIR1, VSIG4), as well as non-immune tissue sources, including neural stem/precursor and meningeal cells (CKB) (Figure 4e). Cell-of-origin enrichment for SIGLEC5 is not shown, as correlated proteins were found across diverse cell types and tissues. Longitudinal analysis demonstrated that these four proteins were markedly suppressed in complete responders, reaching levels comparable to those of HCs by visit 3, whereas non-responders showed minimal change (Figure 4f-j).

## Discussion

In this study, we generated a comprehensive serum proteomic atlas of LN by profiling more than 5,000 proteins in 751 serum samples from 270 LN patients in the longitudinal AMP RA/SLE cohort, revealing extensive molecular heterogeneity underlying the disease. Our analysis uncovered profound systemic proteomic dysregulation, with 2,539 proteins significantly increased in LN patients compared to HCs, a depth of molecular characterization that extends beyond prior studies limited to smaller panels of analytes. Of these dysregulated proteins, increases in VSIG4, HAVCR1, CD27, and LAIR1 consistently emerged as top biomarkers across multiple clinical contexts, including distinguishing proliferative from membranous histological classes and correlations with both NIH AI and CI. Furthermore, 3-month changes in these markers were associated with complete treatment response at one year. Together, these findings suggest that serum proteomics are associated with features of intrarenal immunopathology, offering unique insights into the LN-dominant pathogenesis and a potential non-invasive “liquid biopsy” to refine classification and guide LN management.

A comprehensive proteomic assay represents a powerful advancement in biomarker discovery by extending analysis beyond traditional secreted molecules to a wide array of tissue-derived proteins (27). Unlike conventional assays that measure only circulating cytokines and chemokines, many of the proteins identified here are typically membrane-bound or intracellular, and their aberrant presence in circulation reflects tissue-specific events. For instance, we detected soluble forms of receptors that are actively shed into the circulation via specific proteolytic cleavage events during immune activation (VSIG4 (28), CD27 (29), and LAIR1 (30)) or following epithelial tissue injury (HAVCR1 (31)). Consistent with a role for proteolytic shedding, several matrix metalloproteinases and related enzymes, including MMP3, MMP7, MMP10, and ADAMTS1, were also increased in LN, in line with prior evidence implicating MMP family members in renal injury and remodeling in LN (32, 33). Future studies should validate the contribution of these and other proteases to receptor shedding in LN and define the specific protease-substrate relationships in renal and systemic compartments. Together, these observations suggest that the elevation of these markers in serum may reflect intrarenal pathology, whereas cytokines exhibit rapid, transient fluctuations and lack organ specificity due to their systemic roles in physiology.

Leveraging this ability to detect tissue-specific pathology in circulation, we first examined the molecular signatures underlying current histological classification systems. We found that the histological classification of LN is underpinned by broad molecular differences that extend beyond glomerular morphology. We observed that pure proliferative LN is driven predominantly by inflammatory pathways, including type I IFN and cytokine signaling, consistent with an autoimmune response. In contrast, membranous LN was defined by the dysregulation of cellular adhesion, extracellular matrix organization, and PI3K/AKT signaling. Notably, some analytes contributing to these pathway signatures, such as STAT2 and SMAD family members, are typically intracellular transcriptional regulators; in the serum context, they may be best interpreted as indirect markers of active vesicular trafficking and/or cell turnover rather than sole secreted effectors. Importantly, mixed proliferative and mesangial LN did not form a discrete molecular group but instead spanned the proteomic continuum between these classes, highlighting the limitations of discrete classification systems in capturing the continuous nature of intrarenal pathology. Although current ACR/EULAR guidelines generally do not recommend intensive immunosuppression for mesangial classes (3, 4), these molecular overlaps raise the possibility that a subset of mesangial cases may exhibit more aggressive endotypes than suggested by morphology alone, highlighting the importance of soluble proteomics in potential therapeutic guidance and dissecting understudied pathological mechanisms.

Beyond classification, distinct proteomic signatures associated with NIH AI and CI indices revealed partially divergent biology. Higher AI scores were associated with enrichment of cytokine-mediated signaling and efferocytosis pathways, with proteins such as VSIG4 (tissue-resident macrophage complement receptor (28)), CKB (nervous system and high metabolic demand tissues (34)), MERTK (macrophage tyrosine kinase receptor mediating efferocytosis (35)), and the IL-33 receptor IL1RL1, suggesting that histological activity primarily reflects the influx of activated immune cells and the upregulation of counter-regulatory immune checkpoints. In contrast, higher CI scores were associated with markers such as COL15A1 and the matrix-degrading protease MMP7, which may reflect ongoing structural remodeling. Specifically, COL15A1 functions to anchor the basement membrane to the underlying connective tissue (36); thus, its serum accumulation may indicate the permanent scarring and fusion of the tubulointerstitial compartment that defines the fibrotic end-state. SCF (c-Kit ligand) was also associated with higher NIH CI in our cohort, aligning with its known role in tissue remodeling and renal fibrosis (37), and reproducing our prior observation of SCF as a chronicity-associated biomarker in this same cohort using a Luminex-based assay (17). Additionally, we found complement/coagulation pathways exclusively associated with NIH CI, consistent with a recent large-scale urine proteomic analysis (22) and with data implicating sustained complement activation in tubulointerstitial inflammation and myofibroblast-related fibrosis (38). Together, these findings support a model in which histologic activity reflects inflammatory responses with localized innate immune activity, whereas chronicity captures the downstream remodeling and fibrosis that arise from persistent injury and complement-driven maladaptive repair.

Furthermore, our pathway analysis revealed that cytokine signaling, typically a hallmark of activity, was also significantly associated with chronicity scores. This overlap parallels a recent study that identified active macrophage and T cell infiltrates persisting within scarred tissue, which are associated with a higher risk of long-term kidney failure (39). Similarly, we identified markers that were positively associated with both activity and chronicity, including LAIR1, HAVCR1 (KIM-1), as well as VSIG4, CD27, and CKB. Consistent with the role of CD27 as a costimulatory molecule on activated T and B cells (40, 41), single-cell RNA sequencing of LN kidneys from the AMP network demonstrated local activation of intrarenal B cells with progressive induction of *CD27* and loss of *IGHD/IGHM*. In parallel, the same dataset identified a population of M2-like macrophages with high VSIG4 expression, supporting CD27 and VSIG4 as readouts of renal-resident immune activation rather than a systemic signal (42). HAVCR1 (a known marker of tubular injury (43)) elevation in serum suggests that persistent epithelial damage is shared between acute inflammation and progressive fibrosis. LAIR1, an immune inhibitory receptor that binds collagen (44), likely marks the physical interface where infiltrating immune cells interact with the expanding fibrotic matrix. The soluble form of LAIR-1 may also actively drive progression by antagonizing the inhibitory collagen-LAIR1 interaction (45). Importantly, we found that early reductions in this combined module of inflammation- and injury-associated markers (VSIG4, CD27, CKB, LAIR1, HAVCR1) were strongly predictive of a complete clinical response at 1 year, suggesting that attenuating acute inflammation and infiltrates within fibrotic tissue may be important for preventing long-term failure.

While prior studies have identified associations between select serum and urine soluble biomarkers and LN pathology, these efforts were largely focused on targeted panels measuring fewer than 100 analytes. Collectively, these studies provided critical validation for individual markers identified in this study. In cross-study comparisons, 10 of 14 analyzable proteins (71%) reported by Tang et al. as associated with LN and 26 of 33 transcripts (79%) from the 93-gene SLE MetaSignature described by Haynes et al. were also associated with LN in our cohort, despite the latter being derived from SLE rather than biopsy-confirmed LN (Supplemental Tables 7-8) (46, 47). These concordant findings indicate that our unbiased serum proteomic atlas both recapitulates and extends prior molecular signatures of SLE and LN. For example, TNFRSF1B (TNFR2) emerged as the most important discriminator between LN and healthy controls in our global classifier and was among the top predictors in our preliminary cohort, consistent with the findings of Tang et al. and the role of TNF-TNFR2-dependent endothelial activation and monocyte-to-macrophage transition within inflamed glomerular capillaries^47^ (48). Of particular interest, VSIG4 has been established as a robust serum diagnostic marker (AUC 0.93) that correlates with crescent formation (19, 47). Furthermore, recent analysis of human LN biopsies demonstrated significant colocalization of VSIG4 with IgG and C3, suggesting that VSIG4 may be associated with immune complexes within the kidney and directly contribute to intrarenal pathogenesis (49). Single-cell RNA sequencing of LN kidneys further validates this cellular origin, identifying a specific population of infiltrating M2-like macrophages that highly express VSIG4 and CD163 (42). This provides a mechanistic link to previous large-scale studies that validated urinary CD163 (a macrophage activation marker) as a key predictor of activity and 1-year outcomes (24). The fact that VSIG4 and CD163 mark the same pathogenic cell population supports the concept that biofluids accurately reflect tissue pathology; indeed, transcriptomic profiles of immune cells obtained from urine correlate strongly with those in the kidney (42). Thus, our identification of serum VSIG4 suggests that serum proteomics captures the systemic counterpart to these urinary signals. Notably, although we detected elevated serum CD163 levels in LN, they were not associated with histologic activity, suggesting that CD163 may function predominantly as a urine-enriched readout of broader intrarenal macrophage activation, whereas serum VSIG4 robustly reflects systemic involvement of the same population. Together, these data support a model in which urine provides a sensitive tissue-specific inflammatory readout, whereas serum offers a complementary systemic overview into the same pathogenic cell populations at different tissues.

Furthermore, our data identified additional markers that are traditionally monitored in urine. For example, HAVCR1 and LAIR1 are elevated in the urine of patients with LN, and HAVCR1 correlates with histologic and clinical activity (50–56). Our data further demonstrates their predictive value in serum, offering a unified systemic readout of both immune activation and tissue damage. Importantly, by profiling over 5,000 proteins, our approach significantly extends these observations, uncovering thousands of novel dysregulated proteins and identifying specific isoforms (e.g., HACVR1 vs. HACVR2) associated with distinct histological features. This provides a more granular view of the intrarenal environment, demonstrating a complex, structured interplay among distinct immune and tissue-remodeling molecular modules.

Currently, clinical decision-making is hampered by discordance between standard laboratory values (e.g., proteinuria, creatinine) and the underlying histology. Our data suggest that a targeted panel of serum proteins may be associated with intrarenal pathology. Specifically, distinguishing “active” inflammation (VSIG4, CD27) from “bridging” tissue injury (HAVCR1, LAIR1) could help clinicians distinguish between ongoing immunologic activity requiring intensified immunosuppression and the accrual of chronic damage that may warrant focused renoprotective strategies. Furthermore, the early suppression of a subset of biomarkers, including the VSIG4/CD27/LAIR1/CKB module, in complete responders suggests these biomarkers could serve as early measures of treatment efficacy, identifying non-responders months before conventional clinical criteria would confirm treatment failure. This would enable earlier therapeutic optimization, potentially preventing irreversible damage in patients who are biologically refractory to standard-of-care therapies. Translating these findings onto multiplex platforms suitable for clinical use, such as highly sensitive electrochemiluminescence-based assays, will be an important next step toward implementing these panels in practice.

Our study has several limitations. First, although our cohort of 270 patients is large for a high-dimensional proteomic study, SLE is a highly heterogeneous disease; therefore, validation in independent external cohorts is necessary to confirm the generalizability of these predictive signatures. Second, consistent with the observational design, participants received diverse standard-of-care regimens rather than the standardized protocols used in clinical trials. Third, although serum proteomics provides a window into systemic and localized biology, it remains an indirect measure of tissue pathology. While we identified proteins with known tissue-restricted expression patterns (e.g., VSIG4 in macrophages), paired single-cell RNA sequencing of renal tissue would be required to confirm the cellular source of these circulating analytes. Fourth, our comparison was limited to demographically matched HCs, as we did not include disease controls with non-renal SLE or other glomerulonephritides; further studies are needed to determine which components of these modules are specific to LN. Finally, while our 1-year follow-up was sufficient to assess clinical response, long-term longitudinal tracking is required to determine whether these markers ultimately predict progression to end-stage kidney disease.

In summary, this study presents a comprehensive serum proteomic atlas of LN, revealing distinct molecular signatures that characterize histological activity, chronicity, and treatment response. We identified a core set of biomarkers, including VSIG4, CD27, HAVCR1, and LAIR1, that are associated with activity alone or with activity and chronicity. Importantly, early suppression of these proteins is associated with long-term treatment outcomes, offering a promising tool for precision treatment. At this stage, these markers should be viewed as mechanistic candidates that define molecular endotypes of LN rather than as a fully validated clinical assay, which will require formal biomarker-development studies in additional independent cohorts. These findings underscore the potential of serum proteomics to molecularly inform monitoring and personalized therapeutic management in LN.

## Methods

### Sex as a biological variable

Both male and female participants were included in the HC and SLE cohorts, and the sex distribution did not differ between diagnostic groups. Sex was recorded at enrollment and was included as a covariate in multivariable models.

### Discovery cohort

An independent preliminary cohort of SLE patients (n=27) was recruited from the Oklahoma Rheumatic Disease Cohort, and HCs (n=16) were recruited from the Oklahoma Rheumatic Disease Research Cores Center at the Oklahoma Medical Research Foundation. SLE classification was based on ACR criteria, and renal involvement was defined by meeting the ACR renal criterion. Demographic, clinical, and laboratory data were collected at enrollment, and blood was drawn, processed, and serum stored at −80 °C.

### Confirmatory cohort

Individuals aged 16 years and older diagnosed with SLE (according to the American College of Rheumatology (57) or Systemic Lupus International Collaborating Clinics criteria (58)) who required a kidney biopsy (UPCR ≥0.5) were enrolled at 15 clinical centers participating in Phases 1 and 2 of the Accelerating Medicines Partnership (AMP) RA/SLE Network (59). For the present study, we analyzed a subset of AMP participants for whom baseline serum was available and passed quality control for Olink proteomic profiling. Renal tissue samples were reviewed by a renal pathologist at each site using the ISN/RPS classification system (2) and the NIH activity and chronicity indices (60). The majority were also reviewed and adjudicated by a consensus panel. Exclusion criteria included prior renal transplantation, rituximab treatment within 6 months of biopsy, or pregnancy at the time of biopsy. Participants received standard-of-care therapies as determined by their physicians. The healthy control (HC) group consisted of healthy individuals matched for age, sex, and race/ethnicity, recruited from the Oklahoma Rheumatic Disease Research Cores Center at the Oklahoma Medical Research Foundation and the New York University Langone Health Rheumatology Center.

### Clinical data and sample collection

Information on participant demographics, medical history, and laboratory results was obtained within the two weeks preceding the kidney biopsy. Each site collected laboratory data from local clinical research teams. Peripheral blood specimens were drawn at the time of biopsy (baseline, V0) and at 3 (V1), 6 (V2), and 12 (V3) months post-biopsy. Blood samples were processed and stored at −80°C in accordance with AMP RA/SLE protocol guidelines (59).

### Assessment of clinical response

Clinical outcomes at 12 months were evaluated using the response criteria from the ACCESS trial (61, 62). A complete response was defined by a UPCR ≤0.5, serum creatinine within normal limits (≤1.3 mg/dL) or, if elevated, no more than 125% of baseline value, and corticosteroid therapy reduced to ≤10 mg/day. Partial response was defined as>50% reduction in UPCR without meeting the UPCR criterion for complete response, normal creatinine (≤1.3 mg/DL) or, if abnormal, ≤125% of baseline, and prednisone dose ≤15 mg/day. Non-responders did not meet either set of criteria. Only complete and non-responders were included in the clinical response analyses.

### Olink Assays and Quality Control Process

Serum proteomic profiling was performed using the Olink Proximity Extension Assay (PEA) across the two independent cohorts. For the discovery cohort, protein quantification was performed using the Olink Target 1536 panel, with quality control and normalization performed according to the manufacturer’s standard pipeline, enabling measurement of approximately 1,500 proteins. For the confirmatory (main) cohort, proteomic profiling was performed on the Olink Explore HT platform, which extends the PEA technology to quantify approximately 5,400 proteins, as previously described (27). Briefly, 5 μL of serum per sample was processed according to the manufacturer’s protocol. Protein abundance values were converted to log-transformed protein expression (Log_2_ProtExp) values and scaled by adding a constant of 16 to ensure all values were positive. Transformed values were used for all downstream analyses. Internal assay controls (incubation, extension, amplification) and external controls (sample replicates, plate controls, negative matrix controls) were integrated into each run for technical validation. Quality control steps included: (i) defining lower limits of detection (LLOD) and quantification (LLOQ) from negative matrix controls; (ii) calculating coefficients of variation (CoV) from plate controls, excluding technically variable analytes with CoV >20%; (iii) classifying proteins with >75% of values below the LLOD as categorical, and proteins with >50% of values between LLOD and LLOQ as semi-continuous with censored substitution of LLOQ. For the Explore HT cohort, 1.33% (72/5,421) proteins were excluded during primary QC due to poor technical performance, and an additional 4.75% (254/5,349) were removed because more than 20% of the samples registered values below the LLOD, yielding a final analytic set of 5,095 proteins per sample for downstream analyses.

### Statistics

All primary statistical analyses were conducted using R (v4.1.0) or RStudio (v1.1.463). Normality of continuous variables was evaluated using the Shapiro-Wilk test, and group comparisons were performed using unpaired Student’s t-tests, Wilcoxon rank-sum tests, ANOVA, or Kruskal-Wallis tests as appropriate based on data distribution and the number of comparisons (R, ‘stats’). Adjustments for multiple comparisons utilized the p.adjust function (R, ‘stats’) with the false discovery rate (FDR) correction (Benjamini-Hochberg method). Rank-based correlation analyses were performed using Spearman’s coefficient, with correlation heatmaps generated using the ‘corrplot’ and ‘ggcorrplot’ packages. Cox proportional hazards regression modeling was conducted using time-dependent proteomic and clinical variables (R, ‘survival’).

DEPs were assessed using DESeq2 (v1.32.0), edgeR (v3.34.1), or limma (v3.60.6) in R for prospective regression analyses, adjusting for covariates including age, sex, and genetic ancestry. Proteins with a multi-testing corrected *P* value <0.05 were considered for subsequent analyses. Functional pathway enrichment was performed using gene-set enrichment analysis (GSEA) with clusterProfiler (v4.12.6). Pathways with Padj < 0.05 were considered significant.

Predictive modeling and feature selection were determined using the XGBoost package (v1.7.8.1) in R. For each cross-validation iteration, 75% of the data was randomly assigned as the training set, with the remaining 25% used for testing. This process was repeated across 100 cross-validation runs, and model performance was summarized by mean accuracy and area under the curve (AUC). Receiver operating characteristic (ROC) curves and confusion matrices were constructed using the ‘pROC’ and ‘caret’ packages in R, and 500-fold cross-validation was used for final model validation. Feature importance for each variable was determined by averaging the importance scores from the fitted XGBoost models across the cross-validation runs, and top predictors were defined as those above the elbow point in the ranked importance values.

For dimensionality reduction and data visualization, Uniform Manifold Approximation and Projection (UMAP) projections were performed using the umap package (v0.2.10.0). Cell-type enrichment was assessed using markers and sensitivity data from PanglaoDB. For each analyte, a cell-type enrichment score (CES) was calculated with the formula:

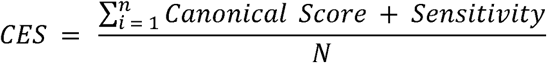

where *N* is the number of proteins. Statistical significance was determined by comparing observed CES values to empirical null distributions generated via 5,000 randomly sampled protein sets. Data manipulation and figure generation employed dplyr, ggplot2, and eulerr, and network or clustering analyses utilized clusterProfiler, dbscan, and iGraph, as appropriate. For all statistical analyses, a *P* or *Padj* <0.05 was considered significant.

### Study approval

All participants provided written informed consent before enrollment, and all procedures were conducted in accordance with protocols approved by the institutional review boards of each participating institution, in compliance with the Declaration of Helsinki.

## Data availability

The data supporting the findings of this study will be deposited in Synapse and made publicly available upon publication.

## Author contributions

R.L., J.M.G, and J.A.J. conceived and designed the study. A.F., P.I., H. M.B., R.M.C., J.A., J.L.B., C.P., D.W., M.H.W, A.D., D.M.F, V.M.H, P.J.U., B.D., J.B., M.P., J.M.G., and J.A.J. provided study materials and/or patients. A.F., P.I., H. M.B., R.M.C., J.A., J.L.B., C.P., D.W., M.H.W, A.D., D.M.F, V.M.H, P.J.U., B.D., J.B., M.P., J.M.G., and J.A.J. collected and assembled data. Data analysis and interpretation: R.L., C.A.W., and B.J. analyzed and interpreted data. Manuscript writing: R.L. and C.A.W wrote the manuscript, and all authors reviewed and edited the manuscript. Accountable for all aspects of the work: R.L., J.M.G and J.A.J. are accountable for all aspects of the work.

