## Supplemental Materials for "Serum proteomic atlas reveals distinct molecular signatures of lupus nephritis activity, chronicity, and treatment response"

**Supplemental Table 1.** Demographics of patients with lupus nephritis (LN) and healthy controls (HCs) in the discovery cohort.

|  | **HCs (n = 16)** | **LN (n=27)** | **p-value** |
| --- | --- | --- | --- |
| **Female, n (%)** | 16 (100) | 27 (100) | >0.9999^1^ |
| **Age, median (IQR)** | 30.5 (24-41) | 29 (24-39) | 0.8277^2^ |
| **Race, n (%)** |  |  | 0.1754^1^ |
| American Indian/Alaskan Native | 1 (6) | 0 (0) |  |
| Asian | 1 (6) | 0 (0) |  |
| Black/African American | 4 (25) | 10 (37) |  |
| White/Caucasian | 7 (44) | 16 (59) |  |
| More than one race | 2 (13) | 1 (4) |  |
| Unknown | 1 (6.3) | 0 (0) |  |
| **Ethnicity, n (%)** |  |  | 0.0133^1^ |
| Hispanic or Latino | 1 (6) | 11 (41) |  |
| Unknown | 0 (0) | 2 (7) |  |
| ^1^Fisher’s exact test; ^2^Mann-Whitney test | | | |

**Supplemental Table 2.** Top predictors for distinguishing healthy controls (n=16) from lupus nephritis (n=27) in the discovery cohort.

|  | **Importance** |
| --- | --- |
| **HAVCR2** | 76.35571 |
| **BST2** | 32.72832 |
| **TNFRSF8** | 21.15914 |
| **CXCL9** | 16.65352 |
| **CD300C** | 14.28281 |
| **CD300E** | 10.06609 |
| **VSIG4** | 8.910005 |
| **VCAM1** | 6.56433 |
| **TNFRSF1B** | 5.564516 |
| **CD27** | 5.255191 |

**Supplemental Table 3.** Top predictors for distinguishing healthy controls (n=64) from lupus nephritis (n=270).

|  | **Importance** |
| --- | --- |
| **TNFRSF1B*** | 81.9666 |
| **VSIG4*** | 41.86422 |
| **HAVCR2*** | 40.61346 |
| **BTN3A2** | 13.9449 |
| **TNFRSF14** | 11.68051 |
| **KPNA7** | 8.5642 |
| **WHRN** | 8.30377 |
| **FBXL7** | 6.249788 |
| **LGALS9** | 5.962196 |
| **PLIN3** | 3.478473 |
| **FAM83C** | 3.396499 |
| **HAVCR1** | 3.32071 |
| **REN** | 3.284741 |
| **DRC1** | 3.203201 |
| **CD1C** | 3.198213 |
| **CD27*** | 2.551113 |
| **IL15** | 2.333427 |
| **RET** | 2.24407 |
| **EPHA4** | 2.172619 |
| **EGFR** | 2.122797 |
| **TINAGL1** | 2.112209 |
| **IFIT3** | 1.735675 |
| **SIRPB1** | 1.636753 |
| **LTA4H** | 1.54272 |
| **DPP4** | 1.429126 |
| **CD300A** | 1.41527 |
| **IFNGR1** | 1.360267 |
| **TXNDC5** | 1.149194 |
| **ARPP21** | 1.09851 |
| **CHAD** | 1.040041 |
| **CNN2** | 0.922597 |
| **SFRP4** | 0.804818 |
| **FGA** | 0.728401 |
| **VCAN** | 0.721868 |
| **KLRB1** | 0.694982 |
| **BST2*** | 0.653813 |
| **DLL1** | 0.647415 |
| **IL3** | 0.597531 |
| **BTD** | 0.580384 |
| **XG** | 0.553999 |
| **FRMD8** | 0.527046 |
| *Identified as top predictors in the discovery cohort | |

**Supplemental Table 4.** Top predictors for distinguishing proliferative (n=98) from membranous lupus nephritis (n=66).

|  | **Importance** |
| --- | --- |
| **CFP** | 61.46561 |
| **BPIFB1** | 51.44582 |
| **APOBR** | 40.29089 |
| **DEFA1** | 39.81057 |
| **CLEC4E** | 37.94561 |
| **PON1** | 33.91289 |
| **MTERF3** | 29.2895 |
| **GPC2** | 28.23845 |
| **CDH17** | 25.1257 |
| **ALPG** | 24.33506 |
| **PALM3** | 23.59405 |
| **ZMYND11** | 22.7148 |
| **DLEU7** | 22.45064 |
| **L1CAM** | 22.01952 |
| **ITGA11** | 20.61531 |
| **PFKFB1** | 19.76862 |
| **MICA_MICB** | 19.7431 |
| **VSIG4** | 19.52292 |
| **CASP8AP2** | 19.16167 |
| **LMNB2** | 18.89209 |
| **DNER** | 18.49796 |
| **RAB3IL1** | 17.57489 |
| **CCL17** | 16.95976 |
| **LRRC27** | 14.42242 |
| **KIAA0232** | 13.65745 |
| **SLITRK5** | 12.98056 |
| **PDXDC1** | 12.84676 |
| **ARID5A** | 12.64911 |
| **ALPP** | 12.29424 |
| **FCRL2** | 9.890041 |
| **SPON1** | 1.367981 |
| **PCDH17** | 1.255098 |

**Supplemental Table 5.** Top predictors of NIH Activity Index (AI) scores in lupus nephritis (n=170).

|  | **Importance** |
| --- | --- |
| **VSIG4** | 46.49171 |
| **IL1RL1** | 24.42284 |
| **C17orf50** | 18.15812 |
| **KEL** | 17.11952 |
| **CDH17** | 15.61857 |
| **SERPINA3** | 13.73583 |
| **TFRC** | 12.91909 |
| **TCL1A** | 11.49912 |
| **ECHDC3** | 10.321 |
| **IL23R** | 9.462675 |
| **FGFR4** | 9.248596 |
| **ZPR1** | 9.24411 |
| **IL3RA** | 8.884698 |
| **CNTN1** | 8.48092 |
| **PGLYRP3** | 8.049561 |
| **MATN3** | 7.743835 |
| **SLC39A14** | 7.559638 |
| **BEND5** | 7.417954 |
| **ARHGAP31** | 7.351263 |
| **ICAM4** | 7.320562 |
| **FCRL5** | 6.898095 |
| **UBXN11** | 6.472559 |
| **ACOT12** | 6.305793 |
| **DPP4** | 5.974443 |
| **CR2** | 5.931836 |
| **C1R** | 5.877558 |
| **SSC4D** | 5.602449 |
| **TINAGL1** | 4.953678 |
| **ACP6** | 4.726832 |
| **DEFA1** | 4.627017 |
| **FNDC1** | 4.580354 |
| **MMP3** | 4.520663 |
| **DNAJA1** | 4.454803 |
| **CR1** | 4.401579 |
| **CHAD** | 4.34684 |
| **ORM1** | 4.204396 |
| **DGKB** | 4.166461 |
| **IL17RB** | 3.765072 |
| **CPXM1** | 3.734467 |
| **AREL1** | 3.726859 |
| **CPA2** | 3.252498 |
| **KLK10** | 3.215557 |
| **PRRC1** | 3.104717 |
| **SIGLEC8** | 3.093604 |
| **GAPDH** | 2.812444 |
| **HAVCR2** | 2.752173 |
| **CA12** | 2.711022 |
| **LYPD8** | 2.559279 |
| **PODXL2** | 2.387153 |
| **CILP** | 2.182669 |
| **MEX3C** | 2.111742 |
| **MAP10** | 2.048306 |
| **TMPRSS5** | 1.969384 |
| **ASGR1** | 1.844736 |
| **KLK7** | 1.817582 |
| **CCL23** | 1.725358 |
| **FOLH1** | 1.68508 |
| **COL1A1** | 1.565439 |
| **HAVCR1** | 1.563754 |
| **ANGPTL1** | 1.534497 |
| **TNFSF12** | 1.400382 |
| **PAMR1** | 1.370447 |
| **AOC3** | 1.125784 |
| **ADAMTS1** | 1.100102 |
| **TNFRSF12A** | 0.705432 |

**Supplemental Table 6.** Top predictors of NIH Chronicity Index (CI) scores in lupus nephritis (n=170).

|  | **Importance** |
| --- | --- |
| **CDNF** | 63.53463 |
| **CLMP** | 29.67706 |
| **MMP7** | 22.92919 |
| **TAFA5** | 16.89006 |
| **TNFRSF19** | 14.34738 |
| **COL15A1** | 14.20148 |
| **XG** | 13.75134 |
| **CYTL1** | 13.33581 |
| **CA14** | 12.1683 |
| **SPINK5** | 11.97691 |
| **DSG3** | 11.15966 |
| **MAK** | 9.719081 |
| **APOBEC2** | 8.520511 |
| **BAMBI** | 7.189526 |
| **SCN4B** | 7.017231 |
| **RAB19** | 6.90426 |
| **CST6** | 6.586869 |
| **EIF4ENIF1** | 6.462044 |
| **ESPL1** | 6.216462 |
| **PRR14L** | 5.071324 |
| **AAAS** | 5.015442 |
| **NCCRP1** | 4.919951 |
| **CST1** | 4.753348 |
| **GAK** | 4.629287 |
| **ADAM22** | 4.453132 |
| **ENDOU** | 4.338456 |
| **NPHS1** | 4.095227 |
| **LY6D** | 3.854743 |
| **WDR24** | 3.809336 |
| **DLK1** | 3.807588 |
| **RNF214** | 3.802057 |
| **MTHFD2** | 3.682284 |
| **IGFBP6** | 3.650716 |
| **RBM27** | 3.475477 |
| **CEACAM19** | 3.203348 |
| **ANAPC10** | 2.968511 |
| **REG1A** | 2.782391 |
| **JAM2** | 2.695547 |
| **LZTS1** | 2.68194 |
| **WFDC12** | 2.200894 |
| **BIN1** | 1.875689 |
| **FOXG1** | 1.832344 |
| **SCGB1A1** | 1.775717 |
| **BSG** | 1.710502 |
| **CD99** | 1.672379 |
| **GUCA2A** | 1.529853 |
| **SLURP1** | 1.5184 |
| **CCL15** | 1.405217 |
| **CCN3** | 1.311028 |
| **CD99L2** | 1.28814 |
| **CD300LG** | 1.192834 |
| **GGT5** | 1.068769 |
| **CCL16** | 1.007815 |
| **CCL14** | 0.957704 |
| **CCL19** | 0.887776 |

**Supplemental Table 7.** Comparison of LN-associated serum proteins in Tang et al. and the current study.

| **Protein** | **Tang et al. result*** | **Current study result** |
| --- | --- | --- |
| **LGALS9** | Up in LN | **Up in LN** |
| **VCAM1** | Up in LN | **Up in LN** |
| **IGFBP2** | Up in LN | **Up in LN** |
| **OPN/SPP1** | Up in LN | **Up in LN** |
| **CD40** | Up in LN | **Up in LN** |
| **TNFSF13B/BAFF** | Up in LN | **Up in LN** |
| **ALCAM** | Up in LN | **Up in LN** |
| **PF4** | Up in LN | NS |
| **TFPI** | Up in LN | NS |
| **BST1/CD157** | Up in LN | Down in LN |
| **TNFRSF1B** | Up in LN | **Up in LN** |
| **CRP** | Up in LN | ND |
| **CD14** | Up in LN | **Up in LN** |
| **CD177** | Up in LN | NS |
| **CSTA** | Up in LN | ND |
| **VSIG4** | Up in LN | **Up in LN** |
| ND, not determined (analyte not available or not evaluable in current dataset); NS, not significant. Bold text indicates concordant associations (10/14 available proteins). *Tang C, Tan G, Teymur A, Guo J, Haces-Garcia A, Zhu W, Williams R, Ning J, Saxena R, Wu T. A serum biomarker panel and miniarray detection system for tracking disease activity and flare risk in lupus nephritis. Front Immunol. 2025 May 1;16:1541907. doi: 10.3389/fimmu.2025.1541907. | | |

**Supplemental Table 8.** Overlap of SLE MetaSignature transcripts from Haynes et al. with lupus nephritis-associated proteins in the current serum proteomic study.

|  | **Hayes et al. result*** | **Current study result** |
| --- | --- | --- |
| **CEACAM1** | Up in LN | **Up in LN** |
| **BST2** | Up in LN | **Up in LN** |
| **SERPING1** | Up in LN | **Up in LN** |
| **CD1C** | Down in LN | **Down in LN** |
| **TYMP** | Up in LN | **Up in LN** |
| **ELANE** | Up in LN | **Up in LN** |
| **GBP1** | Up in LN | **Up in LN** |
| **GRN** | Up in LN | **Up in LN** |
| **IFIT1** | Up in LN | **Up in LN** |
| **IFIT3** | Up in LN | **Up in LN** |
| **LGALS3BP** | Up in LN | **Up in LN** |
| **SAT1** | Up in LN | **Up in LN** |
| **SIGLEC1** | Up in LN | **Up in LN** |
| **TAP1** | Up in LN | Down in LN |
| **TCN2** | Up in LN | **Up in LN** |
| **TNFAIP6** | Up in LN | **Up in LN** |
| **TNFSF10** | Up in LN | **Up in LN** |
| **TNFSF13B** | Up in LN | **Up in LN** |
| **LAMP3** | Up in LN | **Up in LN** |
| **LAP3** | Up in LN | **Up in LN** |
| **ZCCHC2** | Up in LN | **Up in LN** |
| **DDX60** | Up in LN | **Up in LN** |
| **ZBP1** | Up in LN | **Up in LN** |
| **NTNG2** | Up in LN | **Up in LN** |
| **HSH2D** | Up in LN | **Up in LN** |
| **REM2** | Up in LN | **Up in LN** |
| **SAMD9L** | Up in LN | **Up in LN** |
| **RSAD2** | Up in LN | NS |
| **IFIH1** | Up in LN | NS |
| **RTP4** | Up in LN | NS |
| **SAMD9** | Up in LN | NS |
| **FBL** | Down in LN | NS |
| **KLRB1** | Down in LN | NS |
| Of the 93 transcripts in Hayes et al, 32 had corresponding proteins measured in our serum proteomic dataset. NS, not significant. Bold text indicates concordant associations (26 of 33 available proteins). *Haynes WA, Haddon DJ, Diep VK, Khatri A, Bongen E, Yiu G, Balboni I, Bolen CR, Mao R, Utz PJ, Khatri P. Integrated, multicohort analysis reveals unified signature of systemic lupus erythematosus. JCI Insight. 2020 Feb 27;5(4):e122312. doi: 10.1172/jci.insight.122312. | | |

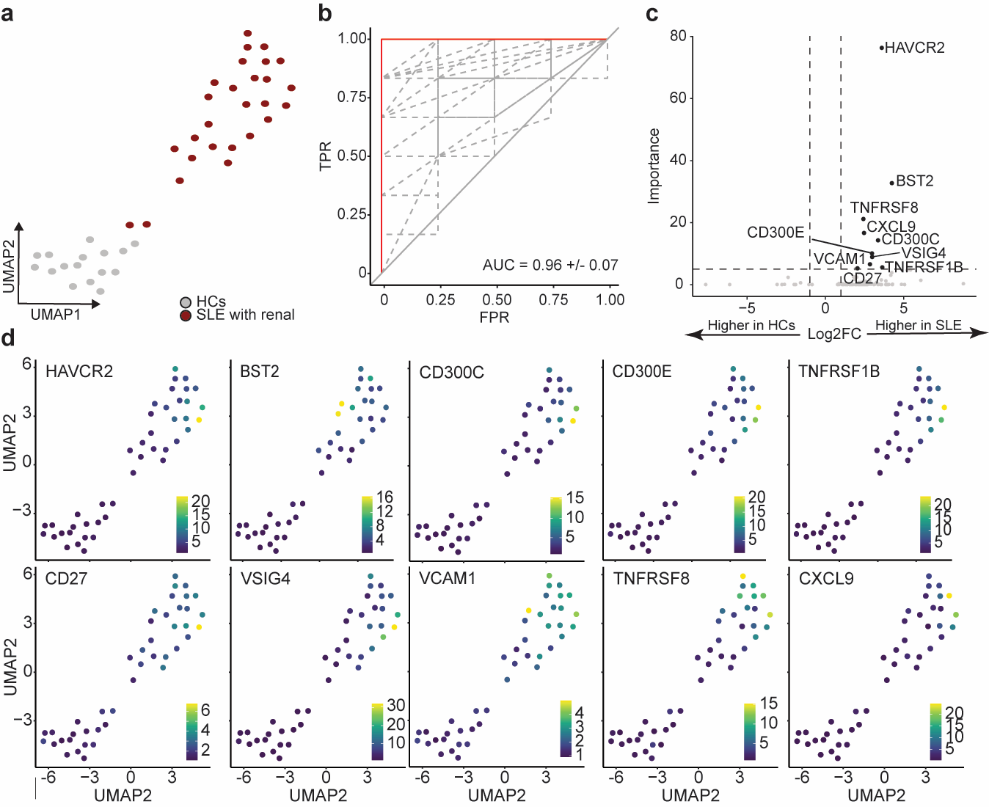

**Supplemental Figure 1. Serum proteins distinguish SLE patients with renal involvement from healthy controls (HCs).** Serum proteins (n=1,536) were quantified using Olink Explore in SLE patients with renal involvement (n=27) and HCs (n=16). **(a)** UMAP plot of all measured serum proteins demonstrates separation between individuals with renal involvement and HCs. **(b)** ROC curve shows the classification performance of an iterative XGBoost model to distinguish SLE with renal involvement from HCs (10-fold cross-validation). Gray dashed lines represent the ROC curves from individual cross-validation iterations, while the red curve denotes the average ROC across all iterations. **(c)** Top 10 protein predictors ranked by SHAP importance and log2 fold change. Dashed lines indicate significance thresholds (*P_adj_* = 0.05) and fold-change cut-offs (>1.2 or <0.8). **(d)** UMAP feature plots display the expression patterns of each top protein predictor. Protein expression values are shown as log2-transformed values.

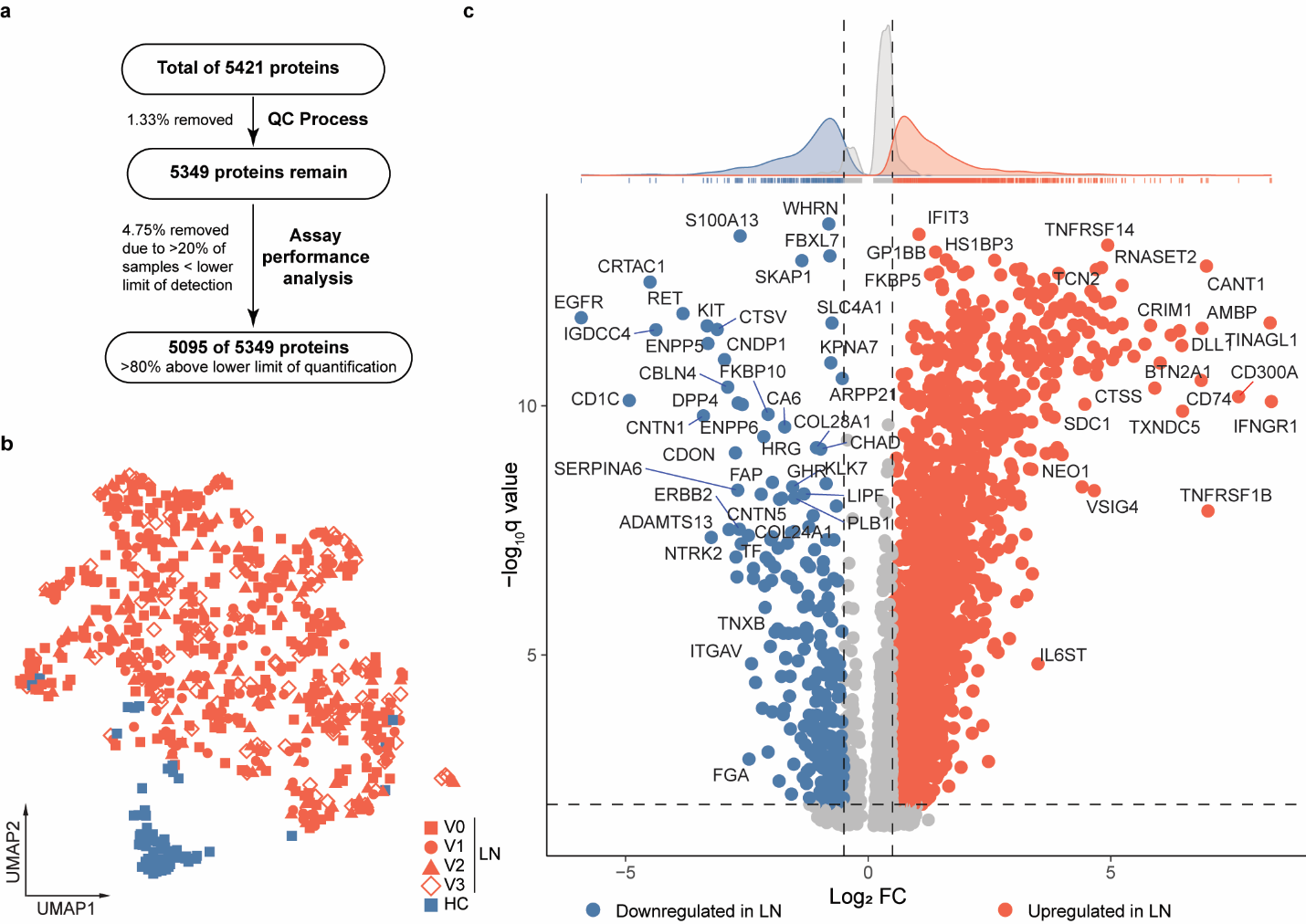
**Supplemental Figure 2. Serum proteomic profile differences between lupus nephritis (LN; n=270) patients and healthy controls (HCs; n=63). (a)** Quality control workflow for Olink Explore HT proteomic data. Starting with 5,421 proteins, sequential filtering removed low-quality measurements (1.33%) and proteins with >20% samples below the lower limit of detection. **(b)** UMAP visualization of longitudinal serum proteomic profiles from LN patients at biopsy (V0; n=270) and follow-up visits at 3 (V1; n=195), 6 (V2; n=159), and 12 (V3; n=127) months post-biopsy compared to healthy controls (HCs; n=64). **(c)** Integrated volcano plot and density distribution of differentially detected proteins between LN patients (baseline) and HCs. Dashed lines indicate significance thresholds (*P_adj_* = 0.05) and fold-change cut-offs (±0.2). Statistical significance was determined using logistic regression, adjusting for covariates including age, sex, and genetic ancestry.

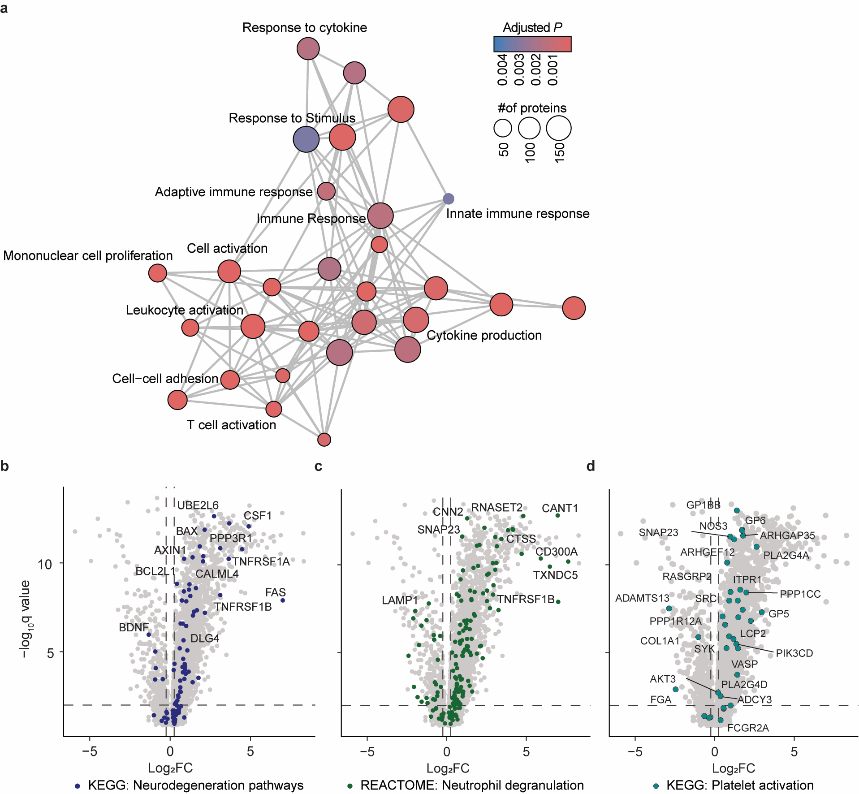

**Supplemental Figure 3. Enriched pathways in lupus nephritis (LN) compared to healthy controls (HCs). (a)** Heatmap of significantly enriched GO pathways in LN (n=270) compared to HCs (n=63). **(b-d)** Volcano plots showing protein enrichment from **(b)** KEGG neurodegeneration, **(c)** REACTOME membrane trafficking, and **(d)** KEGG platelet activation pathways. Dashed lines indicate significance thresholds (*P_adj_* = 0.05) and fold-change cut-offs (±0.2). Statistical significance was determined using logistic regression, adjusting for covariates including age, sex, and genetic ancestry.

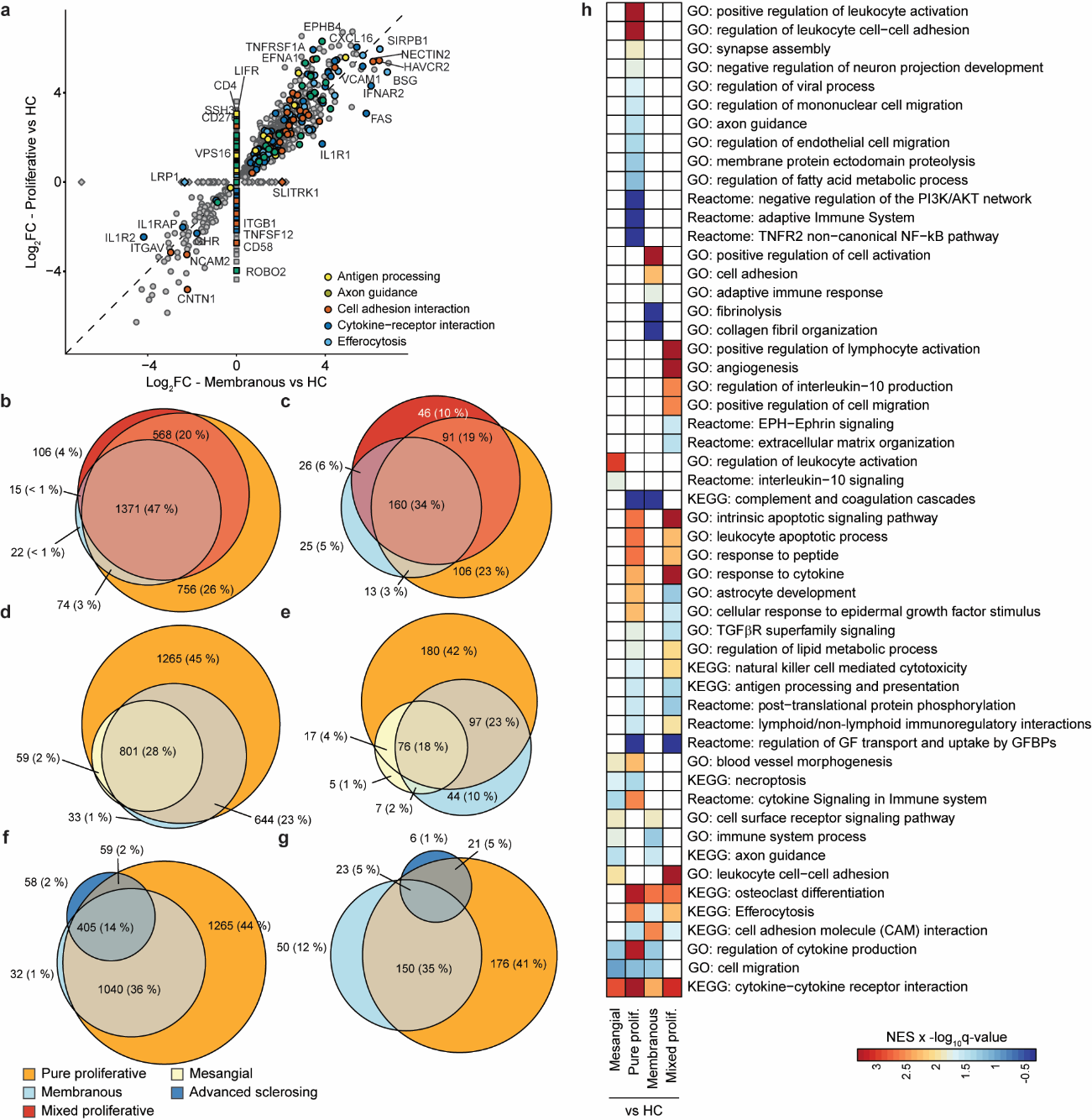

**Supplemental Figure 4. Shared and class-specific proteomic alterations in ISN classes compared to healthy controls. (a)** Scatter plot comparing protein differential expression patterns between pure proliferative LN (class III/IV; n=93) and membranous LN (class V; n=64) relative to healthy controls (HCs). Proteins are colored according to select KEGG pathways. **(b-g)** Venn diagrams illustrating the overlap of DEPs (q<0.05 vs. HCs) among pure proliferative, membranous, and either **(d, e)** mixed proliferative (class III/IV+V; n=78), **(f, g)** mesangial (class I/II; n=23), or **(h, i)** advanced sclerosing (class VI; n=12) LN classes. **(b, d, f)** show upregulated DEPS; **(c, e, g)** show downregulated DEPs relative to healthy controls. **(h)** GSEA pathway enrichment heatmap comparing membranous, pure proliferative, and mixed proliferative LN classes to healthy controls. Color intensity represents the normalized enrichment score (NES) multiplied by the -log_10_q-value. Statistical significance was determined using logistic regression, adjusting for covariates including age, sex, and genetic ancestry.

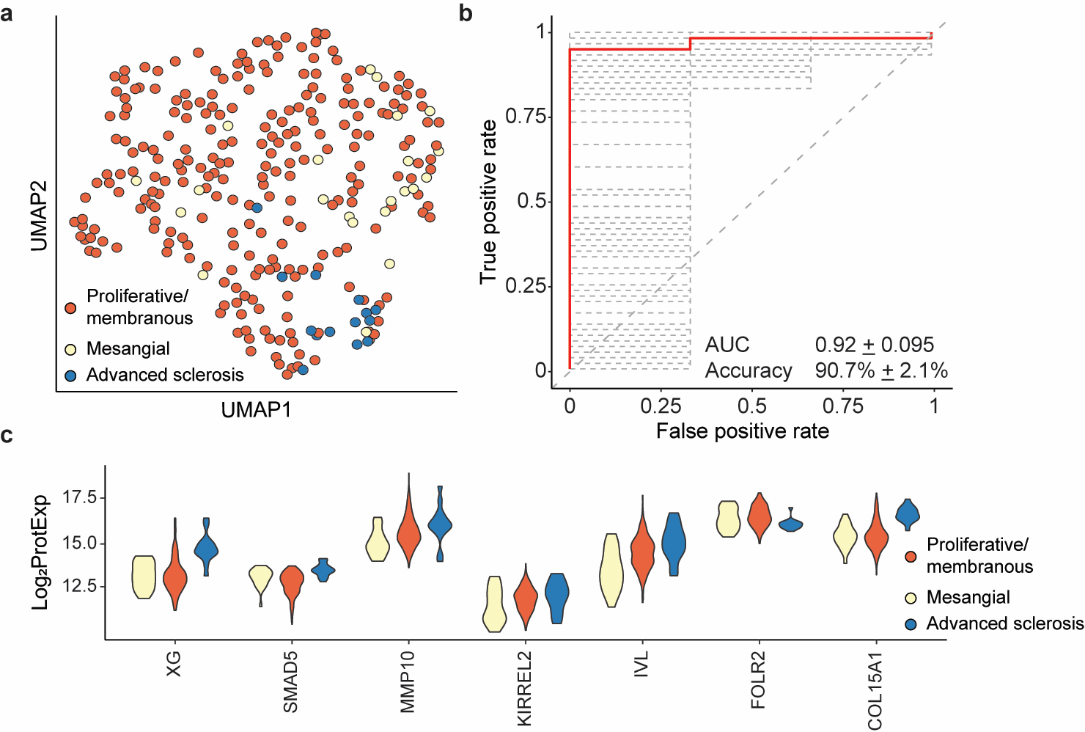

**Supplemental Figure 5. Advanced sclerosing LN classification using machine learning. (a)** UMAP visualization of serum proteomic profiles for advanced sclerosing (class VI; n=12), membranous/proliferative (n=246), and mesangial (class I/II; n=23) LN disease classes, based on the top discriminatory proteins identified by iterative XGBoost analysis. **(b)** ROC curve demonstrating the classification performance of the iterative XGBoost classifier in distinguishing advanced sclerosing LN from other classes, evaluated by 500-fold cross-validation. Gray dashed lines represent the ROC curves from individual cross-validation iterations, while the red curve denotes the average ROC across all iterations. **(c)** Violin plots displaying the expression levels (log_2_-transformed) of the top discriminatory proteins for advanced sclerosing LN across the 3 groups.

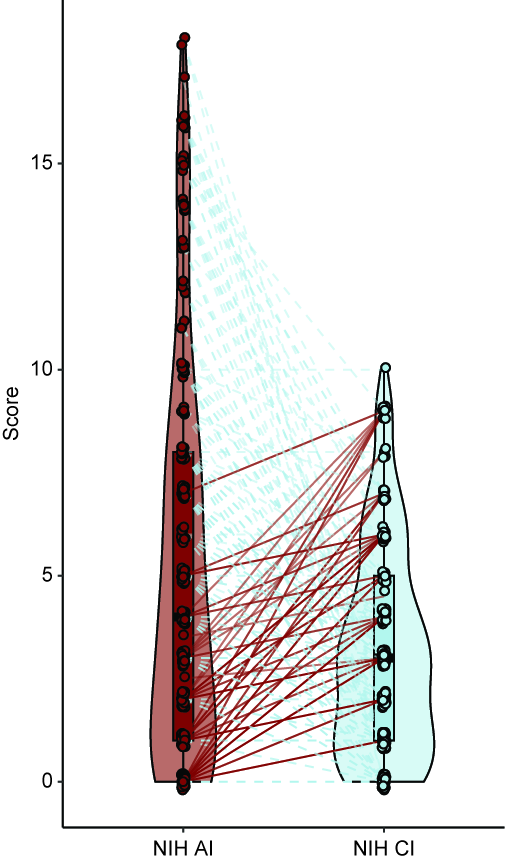

**Supplemental Figure 6. Paired distribution of NIH activity (AI) and chronicity index (CI) scores.** Violin plots show the distribution of NIH AI and CI scores for individual patients (n=170). Each circle represents a single biopsy’s score, and lines connect paired AI and CI scores for the same patient.

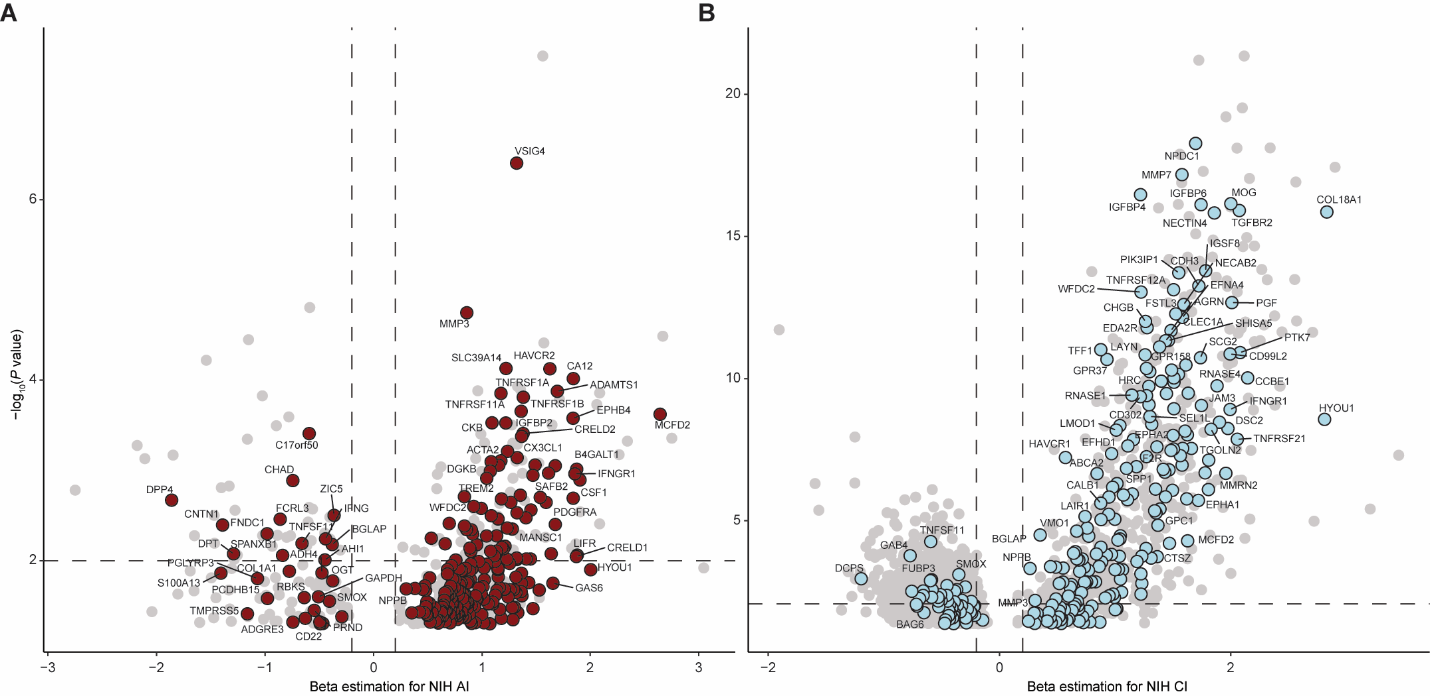
**Supplemental Figure 7. Protein associations with NIH activity (AI) and chronicity indices (CI).** Logistic regression analyses were performed for associations with AI and CI using an iterative decision-tree-based machine learning algorithm. Volcano plots display the beta estimations of individual proteins with the **(a)** AI and **(b)** CI. Statistical significance was determined using linear regression, adjusting for covariates including age, sex, and genetic ancestry.

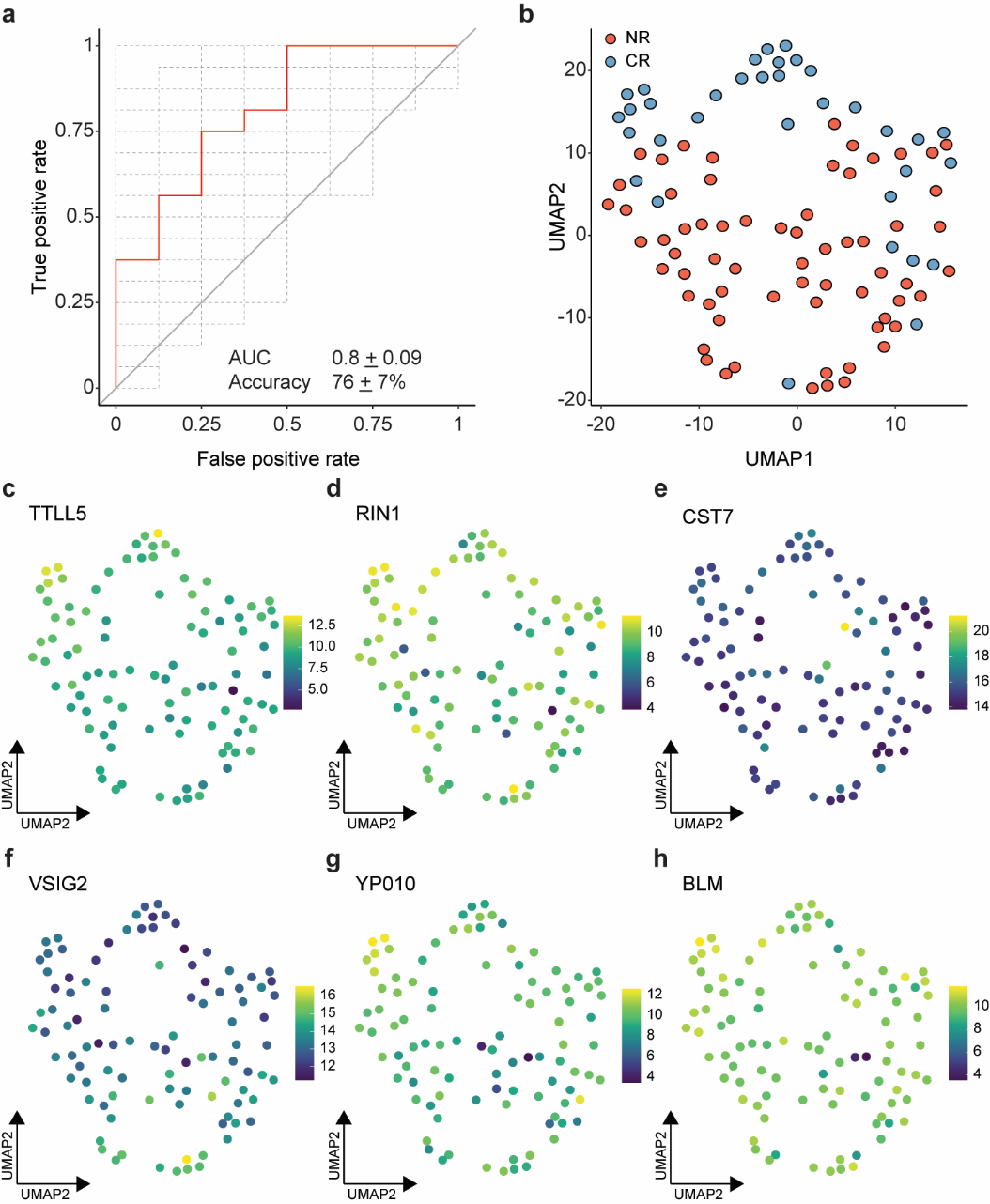

**Supplemental Figure 8. Baseline serum proteomic profiles modestly predict 1-year treatment response in lupus nephritis (LN)** **(a)** Receiver operating characteristic curve showing the performance of the iterative XGBoost classifier in distinguishing patients with complete response (CR; n=32) from those with no response (NR; n=66) at 1 year. The model was initially trained using all available baseline protein levels, and the most discriminatory predictors were selected through an iterative feature elimination process. ROC curves from individual 500-fold cross-validation iterations are shown as gray lines, while the red curve denotes the average performance across all iterations. **(b)** UMAP visualization of serum proteomic profiles for patients with CR and NR, using the top discriminatory proteins. **(c-h)** Feature plots displaying the expression patterns of the top six baseline predictors for treatment response, overlaid on the UMAP embedding. Protein expression values are log_2_-transformed.

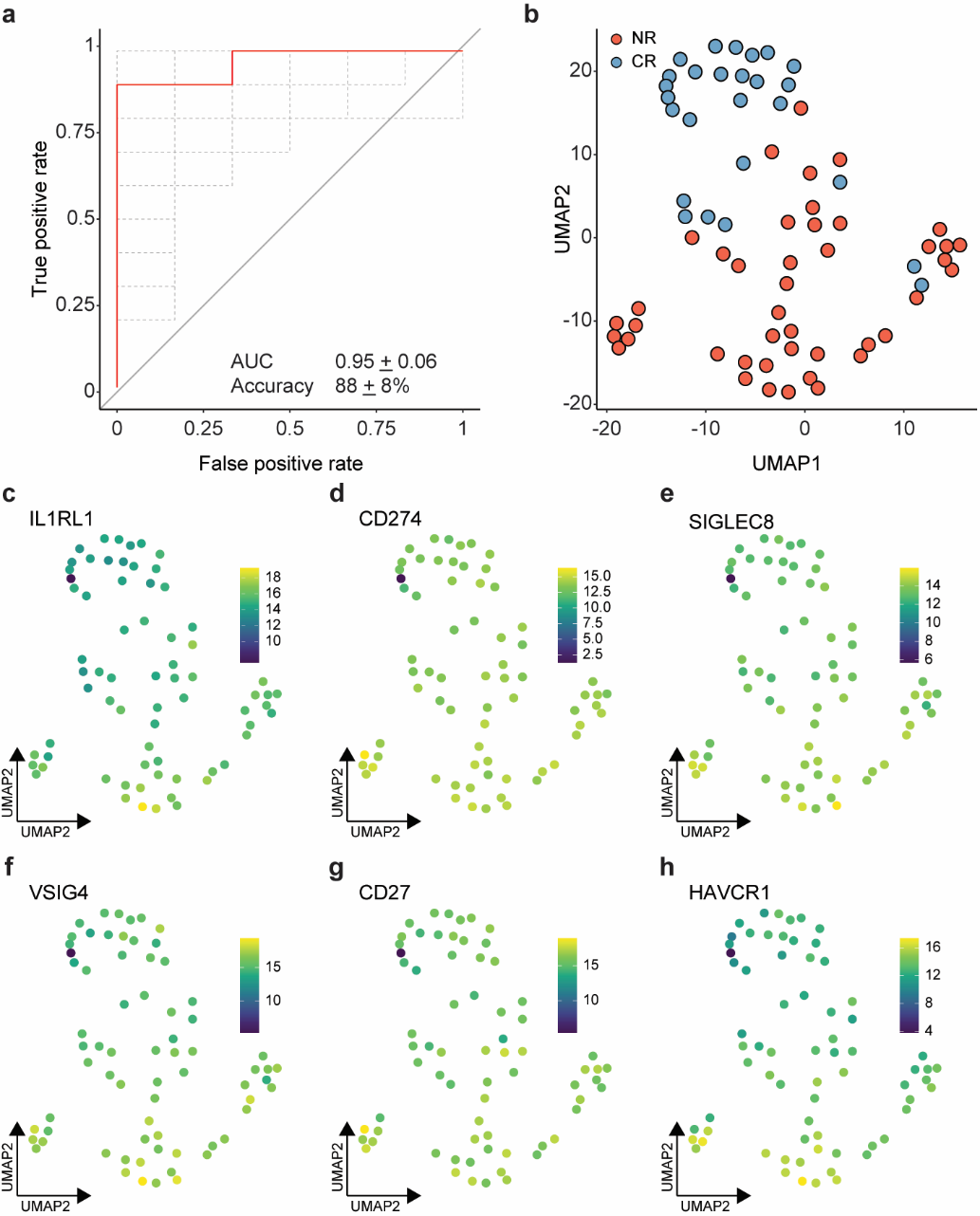

**Supplemental Figure 9. One-year serum proteomic profiles accurately classify 1-year treatment response in lupus nephritis (LN) (a).** Receiver operating characteristic curve showing the performance of the iterative XGBoost classifier in distinguishing patients with complete response (CR; n=32) from those with no response (NR; n=66) at 1 year. The model was initially trained using all available visit 3 (1 year) protein levels, and the most discriminatory predictors were selected through an iterative feature elimination process. ROC curves from individual 500-fold cross-validation iterations are shown as gray lines, while the red curve denotes the average performance across all iterations. **(b)** UMAP visualization of serum proteomic profiles for patients with CR and NR, using the top discriminatory proteins. **(c-h)** Feature plots displaying the expression patterns of the top six baseline predictors for treatment response, overlaid on the UMAP embedding. Protein expression values are log_2_-transformed.

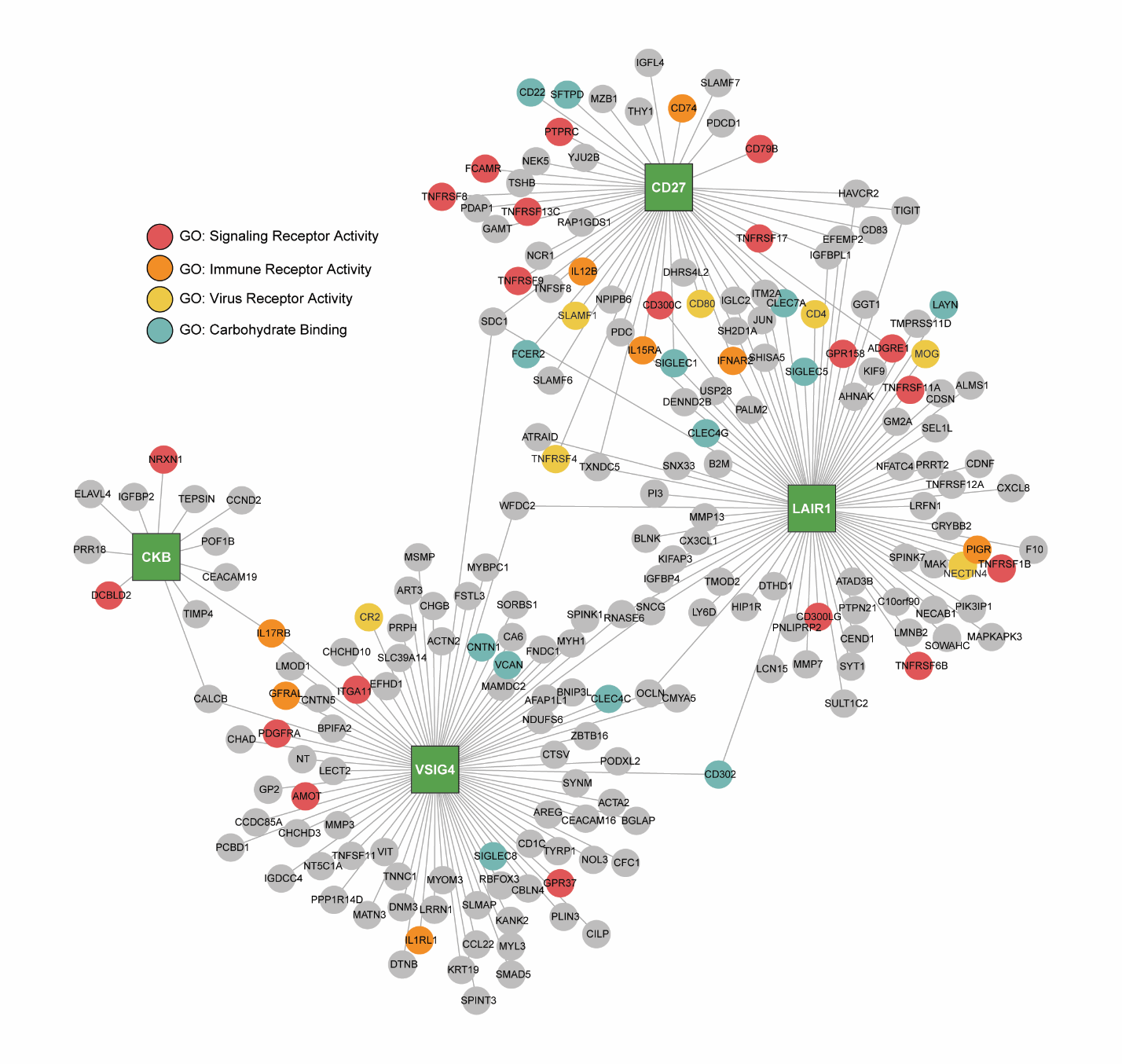

**Supplemental Figure 10. Highly correlated proteins that change at 3 months in patients with a complete response are associated with immune-related surface receptor activity.** Gene regulatory network analyses of the top 4 correlated predictors of treatment response demonstrate that these proteins are centrally positioned within a network enriched for immune-related surface receptor binding and signaling pathways. Nodes represent proteins, with the top 4 predictors emphasized, and edges indicate known or predicted regulatory relationships.
